# State-dependent modulation of cross-area sensory processing in mouse neocortex

**DOI:** 10.64898/2026.10.30.763148

**Authors:** Shuting Han, Fritjof Helmchen

## Abstract

Perception and decision-making rely on distributed cortical circuits that transform sensory inputs into choices and actions. Such cortical processing varies with behavioral state, such that the same inputs can be handled differently across states. However, how behavioral state shapes sensory processing across distributed brain regions remains unclear. To address this question, we trained mice to perform a two-alternative choice task that required discrimination between paired auditory and whisker cues. During task training, we simultaneously recorded neuronal population activity from the primary somatosensory cortex (S1) and the posterior parietal cortex (PPC), using two-photon imaging of neurons expressing the fluorescent calcium indicator GCaMP6f. Using a Hidden Markov Model with Generalized Linear Models (GLM-HMM), we identified three latent behavioral states, including one optimal-performing stimulus-guided state and two sub-optimal biased states. We found that the optimal state was associated with stronger sensory representation in S1 and PPC, as well as stronger bottom-up information transfer from S1 to anterior PPC (PPC-A). In contrast, biased states showed weaker sensory coding and relatively stronger involvement of rostrolateral PPC (PPC-RL). State-dependent differences were present in both task-related and task-orthogonal population activity. These results show that internal states modulate not only local sensory encoding, but also the routing of task-relevant information across cortical networks.

## Introduction

The brain processes external inputs dynamically, adjusting how information is represented and used depending on both external task context and internal behavioral state. A growing body of work demonstrates that distinct behavioral states, such as arousal^1–3^, attention^4,5^, locomotion^6–8^, and motivation^9,10^, profoundly influence sensory responses and bias subsequent decision-making. These state fluctuations occur on both fast and slow timescales and coincide with changes in the activity of individual neurons and in neuronal population variability structure^11–16^. In addition, behavioral state can also alter how task-relevant sensory information is routed across distributed cortical circuits^17–19^. Classic work on the communication-through-coherence theory proposes that inter-area coupling and oscillatory synchrony can gate effective communication between areas^20^, and coordinated inter-area synchrony has been linked to improved task performance^21^. However, we still poorly understand how behavioral states shape the cross-area transfer of task information in distributed brain regions, and how these changes drive state-dependent behavioral output.

Performing a behavioral task requires the brain to transform relevant sensory inputs into corresponding actions. A key area in this sensorimotor transformation is the posterior parietal cortex (PPC). PPC integrates multisensory information across modalities and routes sensory information in a modality– and subregion-dependent way: in mice, the rostrolateral PPC (PPC-RL) routes visual and tactile information, and the anterior PPC (PPC-A) routes visual and auditory information^17,22,23^. PPC also encodes cross-modal prediction and can influence sensory coding and behavioral choices in a state-dependent manner: behavioral choices guided by predictions are associated with faster reaction times and top-down information flow from PPC-A to S1, whereas choices guided by sensory evidence show slower reaction times and bottom-up flow from S1 to PPC-A^22^. Moreover, recent studies show that PPC routes information to different parts of the cortex depending on the movement state^17,18^, and that PPC activity directly reflects behavioral engagement state^24^. This evidence makes PPC a key area for understanding state-dependent sensory processing in the neocortex.

To study state-dependent sensory processing, one challenge lies in inferring internal behavioral states from observable behavior. In recent years, Generalized Linear Model-Hidden Markov Models (GLM-HMMs) have become powerful tools for identifying latent behavioral states that reflect interpretable internal states, providing a useful framework for linking latent behavioral states to observable behavior and brain activity^10,19,25^. Despite the advances in behavioral modeling, it remains unclear how these behavioral states modulate the transformation and communication of task-relevant information across distributed brain regions. Addressing this question requires approaches that capture neural population activity across multiple areas simultaneously and relate it to moment-to-moment changes in behavior and latent state. Here, we combine simultaneous recordings across the cortical hierarchy in a multisensory two-alternative choice paradigm with behavioral modeling to understand how behavioral states shape sensory processing and inter-areal communication. Specifically, we trained mice to perform a task where they learnt to discriminate two combinations of auditory tones and whisker tactile cues. During task training, we simultaneously recorded layer 2/3 (L2/3) neuronal population activity in S1 and subregions of PPC (RL or A) using two-photon calcium imaging with GCaMP6f. Using GLM-HMM, we identified distinct behavioral states that are associated with different levels of local task representation and inter-area communication between S1 and PPC. These results identify PPC as a key node in state-dependent sensory processing and inter-areal routing of sensory information.

## Results

### Behavioral task and two-area two-photon recordings

We trained 16 mice to perform a two-alternative auditory-cued texture discrimination task (Figure 1A). Each trial began with a high– or low-frequency tone for 1 second. Then, a sandpaper (rough or smooth, fixed pairing with the tone) was presented onto the whisker pad of mice for 1 second. Then, following a tone that indicated the start of a choice window, mice chose between two lick ports (left or right) to receive a water reward. The choice window lasted for 2 seconds, and the first lick in this window triggered the outcome window (see Methods). All mice successfully learned the task, completing 100–500 trials per day (Figure 1C). To record task-related neural activity, we used a custom two-area two-photon microscope to simultaneously image from L2/3 neurons of the S1 whisker barrel field and one of the two PPC subregions, PPC-A or PPC-RL (Figure 1B). All mice were transgenic strains expressing the calcium indicator GCaMP6f in excitatory neurons in cortical layer 2/3 (Methods). This dataset has been published in our previous studies where we discovered that the anterior subregion of PPC, PPC-A, is involved in audio-tactile learning and prediction, whereas the rostrolateral subregion PPC-RL is only involved in tactile processing^22,23^. As in our previous study, we defined three learning phases: the naïve phase, where the session behavioral performance was below 55%; the learning phase, where the performance was between 55% and 75%; and the expert phase, where the performance exceeded 75% correct.

**Figure 1.**
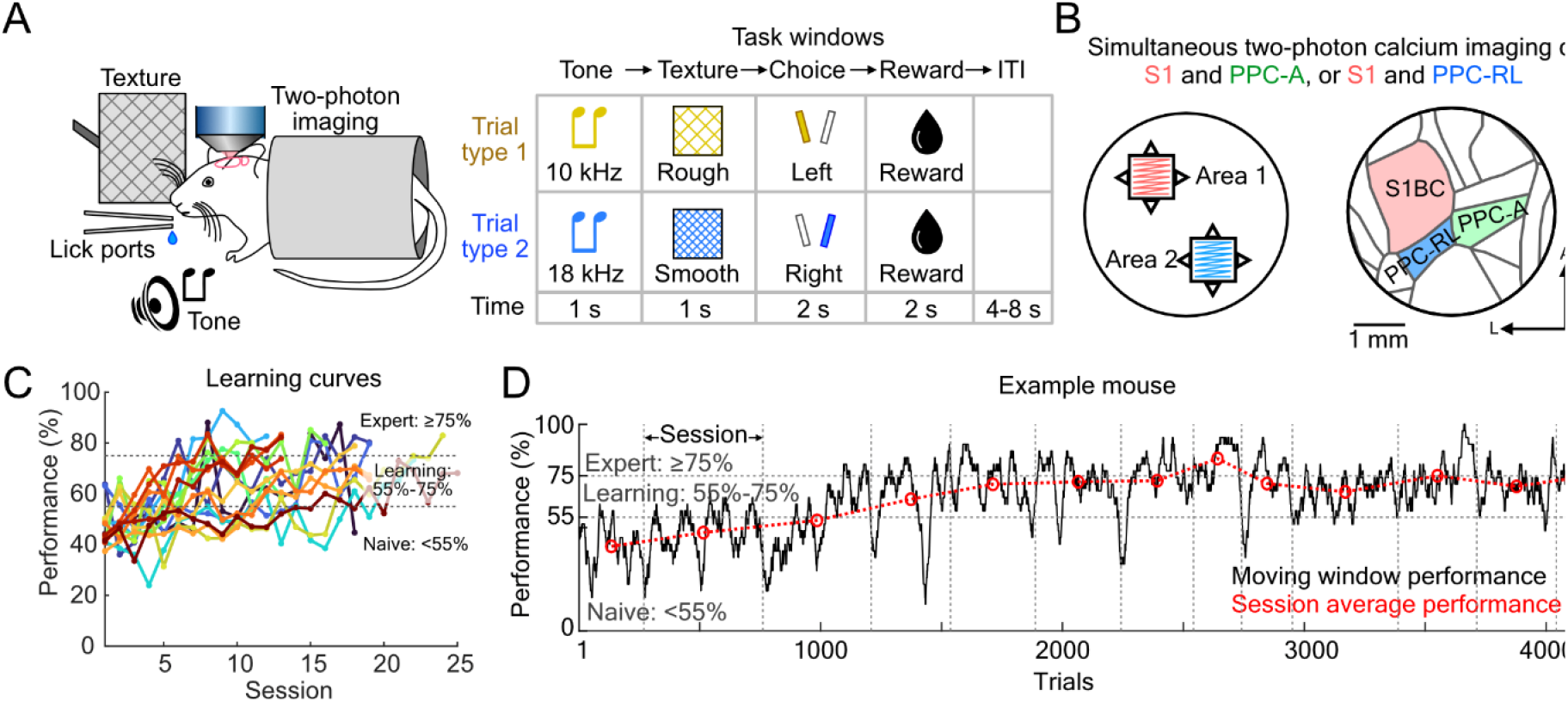
Two-alternative sensory discrimination task. (**A**) Schematic of the experimental setup. Mice were head-fixed under the two-photon microscope, and were presented with an auditory tone followed by a sandpaper texture stimulus. Two lick ports positioned in front of the mice delivered water reward depending on the choice. (**B**) Schematic of two-area two-photon imaging. Simultaneous imaging was performed in S1 barrel cortex (BC) and either of the PPC subregions, PPC-A or PPC-RL. (**C**) Learning curves of all mice. Performance was computed as *N*correct/(*N*correct + *N*incorrect) using all trials within each day, where *N*correct is the total number of correct trials for both trial types, and *N*incorrect is the total number of incorrect trials. (n=16 mice) (**D**) Task performance of an example mouse over all sessions. Vertical dashed lines indicate boundaries between daily training sessions; black line indicates performance using a 30-trial moving window; red circles indicate the session average performance.

As mice learnt the task, we noticed that the overall performance steadily improved over days, but the within-session performance fluctuated, with some blocks of trials exceeding expert performance (>75% success rate) even when the session average remained below expert level (Figure 1D). These fluctuations suggest that mice learnt the task rule earlier than indicated by the sessions’ average performance, but that task performance at this early learning stage remained unstable. One potential cause for the performance fluctuations is variation in behavioral state, which may result in variable task engagement levels. Such behavioral state fluctuations have been reported in many rodent decision-making tasks and may arise from multiple factors, including changes in arousal and attention^10,16,24,26^.

### Identifying behavioral states using GLM-HMM

To identify the behavioral states of mice, we fit Bernoulli GLM-HMMs to predict the behavioral choice of mice^25,27^ (Figure 2A; Methods). We first performed a model selection to identify the most informative predictors, including stimulus (as tone and texture provided the same information in this task), previous choice (lick left or lick right; up to three previous trials), strategy (win-stay-lose-switch), previous reward (up to three previous trials), and a constant term capturing intrinsic bias. Using the behavioral data from all mice, we found that a two-predictor model using stimulus and bias achieved optimal performance with three latent states (Figure 2B-C). These three states corresponded to one optimal-performing, stimulus-driven state (“optimal state”, State 1), in which stimulus information but not bias predicted choice, and two bias-dominated states (“biased state”), in which the bias term but not stimulus information predicted choice (left bias and right bias, States 2 and 3; Figure 2D). Previous studies have discovered similar behavioral states during discrimination tasks^3,10,16,19,24,26,27^, where the states were often termed “engaged state” and “disengaged state”. In our case, because mice continued to make active choices in the left and right choice-biased states, we decided to refer to the states as “optimal state” and “biased states” instead. The previous studies suggest that mice share common behavioral states, although the transition probabilities between these states could vary according to learning phases. To account for this possibility, we fixed the state-specific GLM weights (Figure 2D) and refitted the state transition matrix for each learning phase (Figure 2E; Methods). These learning-specific models performed comparably in prediction accuracy (Figure 2F). As mice progressed from naïve to expert phase, the biased states became less stable, and the optimal state emerged during learning (Figure 2E,G; Figure 2-figure Supplement 1), indicating that task-learning promotes the stabilization of task performance and engagement.

**Figure 2.**
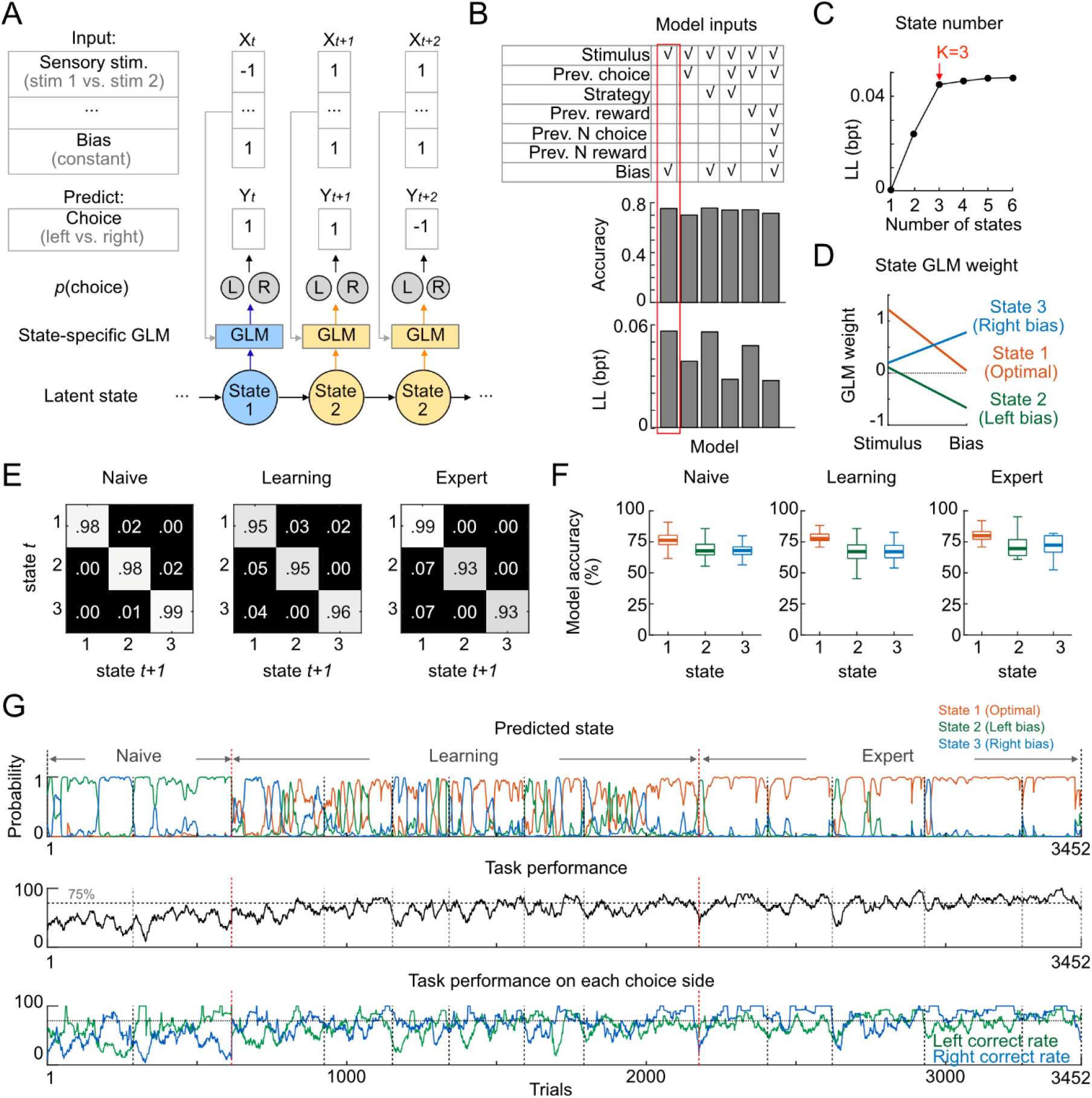
Identification of latent behavioral states using GLM-HMM. (**A**) Schematic of the GLM-HMM design. Task variables were used to predict the behavioral choice of mice through a GLM-coupled HMM. Each latent state is coupled with a state-specific GLM, which predicts the choice probability. (**B**) Top: model comparison using different task variables, including stimulus, previous trial choice, strategy (win-stay-lose-switch), previous trial reward, additional choice history from three previous trials (“Prev. N choice”), additional reward history from three previous trials (“Prev. N reward”), and a constant (“Bias”). Bottom: test set model performance, measured by prediction accuracy and log-likelihood in bits per trial (bpt). The red rectangle highlights the model using stimulus and bias as inputs, which we used for all subsequent analyses. (**C**) Model comparison using different numbers of latent states. Test set performance was measured using log-likelihood (LL) in bits per trial (bpt). The red arrow highlights a three-state model, which we used for all subsequent analyses. (**D**) Inferred GLM weight for the three-state model. State 1 has high weights on the stimulus, indicating stimulus-guided decision and a high-performance state; state 2 and 3 have high weights on the bias term with opposite signs, indicating decision driven by left or right bias. (**E**) Inferred transition matrix using behavior data from separate learning phases, and with the GLM weights fixed as in D. The naïve phase was defined as session performance <55%; the learning phase was defined as performance between 55% and 75%; the expert was defined as performance ≥ 75%. With task-learning, the transition probability from state 2 and 3 to state 1 gradually increased, indicating that the biased states became less stable whereas the optimal states became more robust. (**F**) Model prediction accuracy from each learning phase model. (**G**) Model prediction and task performance from an example mouse. Top: predicted state probability for each state; middle: moving window task performance using a 30-trial window; bottom: task performance for each choice side, for the left or right lick ports. (n=16 mice, 278 sessions; naïve: 117 sessions, learning: 96 sessions, expert: 65 sessions)

We verified the three behavioral states by examining multiple behavioral measurements. First, we compared the overall success rate and the success rate for the left or right lick port in each behavioral state. Mice exhibited high success rate for both lick ports during the optimal state, while the biased states showed a high success rate only for their biased side (Figure 3A). Next, we characterized the state probability within each session and observed learning-related changes. In the naïve phase, mice resided in one of the two biased states throughout the sessions; in the learning phase, mice showed a higher likelihood of being in the optimal state towards the end of the sessions, suggesting within-session learning; in the expert phase, mice quickly entered the optimal state and remained high-performance throughout the sessions (Figure 3B). Finally, we characterized pupil diameter, body movement, face movement, and lick patterns across states (Figure 3C-F). Mice showed larger pupil diameters during the optimal state than during the biased states in both correct and incorrect trials, suggesting that the optimal state captured a condition in which mice exhibited higher arousal level and greater attention to task-relevant inputs (Figure 3C). In contrast, the body and face movement of mice remained at comparable levels across behavioral states (Figure 3D-E). Mice also showed faster decision times during the biased states than during the optimal state, suggesting that choices in these states were more strongly driven by a pre-existing choice bias (Figure 3F). Overall, these results show that the behavioral states identified by GLM-HMM capture differences in task performance and arousal levels. For subsequent analyses, we combined left– and right-bias states into a single “biased” state.

**Figure 3.**
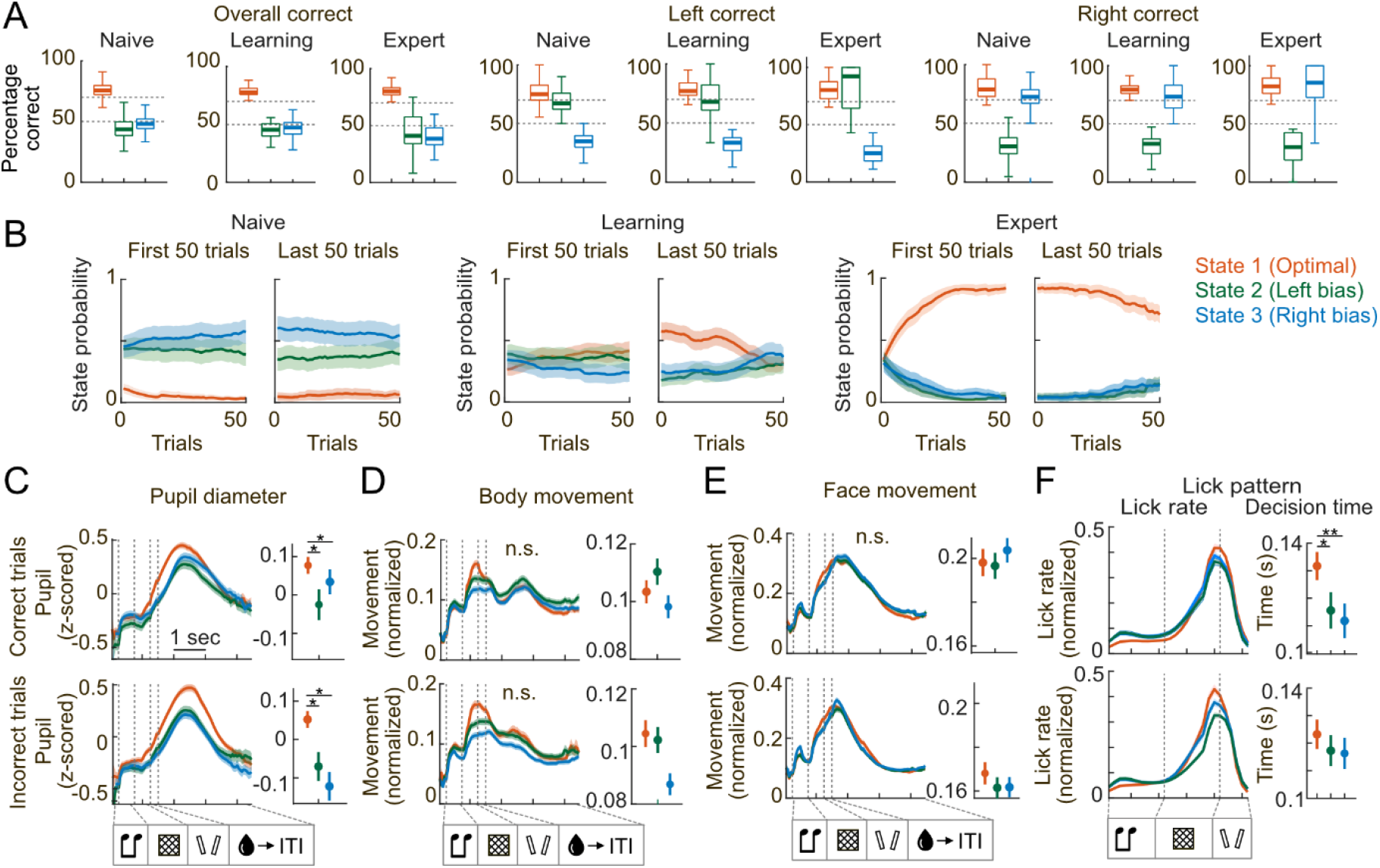
Behavioral characteristics of the identified latent states. (**A**) Task performance for both lick ports combined (“overall correct”), for the left lick port (“left correct”), and for the right lick port (“right correct”). Color indicates state: red represents state 1, green represents state 2, blue represents state 3. State 1 showed high performance on both lick ports, whereas state 2 and 3 only showed high performance on their biased side. (**B**) State probability in the first and last 50 trials for each session. (**C**) Pupil diameter for each state, during correct (top) and incorrect (bottom) trials. Pupil diameter was z-scored within each session. (**D**) Body movement for each state, during correct (top) and incorrect (bottom) trials. Movement was normalized to [0, 1] within each session. (**E**) Face movement for each state, during correct (top) and incorrect (bottom) trials. Movement was normalized to [0, 1] within each session. (**F**) Lick pattern for each state. Left: normalized lick rate in correct (top) and incorrect (bottom) trials. Right: decision time in each state. To ensure stable results, only sessions with more than 20 trials in the states were included for analysis. (n=16 mice, 170 sessions; \**p*<0.05; \*\**p*<0.01)

### Population activity in S1 and PPC reflects behavioral states

To understand whether neocortical sensory processing differs across behavioral states, we investigated the neuronal population activity across S1 and PPC regions. Because mice were not punished for early licks during texture presentation, we removed all time points during which mice licked in the texture window to avoid contamination by lick-related neural activity. In addition, due to the variable choice-window length (mice were free to choose any time during the 2-s choice window), we defined the choice window as the 0.5-s window preceding the lick event that triggered the outcome window (reward or no reward), as in our previous studies^22,23^. Since the optimal state and the biased state contained different numbers of trials and had different correct rates, we only analyzed the correct trials from both states and bootstrapped the trial numbers to be matching across states. We repeated the bootstrapping procedure 20 times for each dataset and averaged the results. These procedures minimized the effect of actions and outcomes linked to different trial types.

We first assessed the amount of task information carried by each population across behavioral states. We trained linear decoders to read out tone, texture, and choice information from their corresponding task windows (Figure 4A; Methods). We found that S1 and PPC-A contained higher levels of texture information during the optimal state than during the biased state, suggesting that task-relevant sensory information was better represented in these populations in the optimal state (Figure 4B). S1 and PPC-A populations also showed slightly higher texture-related activity during the optimal state than during the biased state, although this difference was not significant (Figure 4 – figure supplement 1). We then asked whether population activity in S1 and PPC directly reflected behavioral state. To answer this question, we trained linear decoders to decode the state (optimal vs. biased) from the population activity across task windows (Figure 4C). While behavioral states could be decoded only during the texture window in S1 and PPC-RL, the activity in PPC-A consistently carried information about the behavioral state across task windows (Figure 4D). These findings are consistent with previous studies showing that PPC activity reflects behavioral engagement and choice bias during decision-making tasks^24,28,29^.

**Figure 4.**
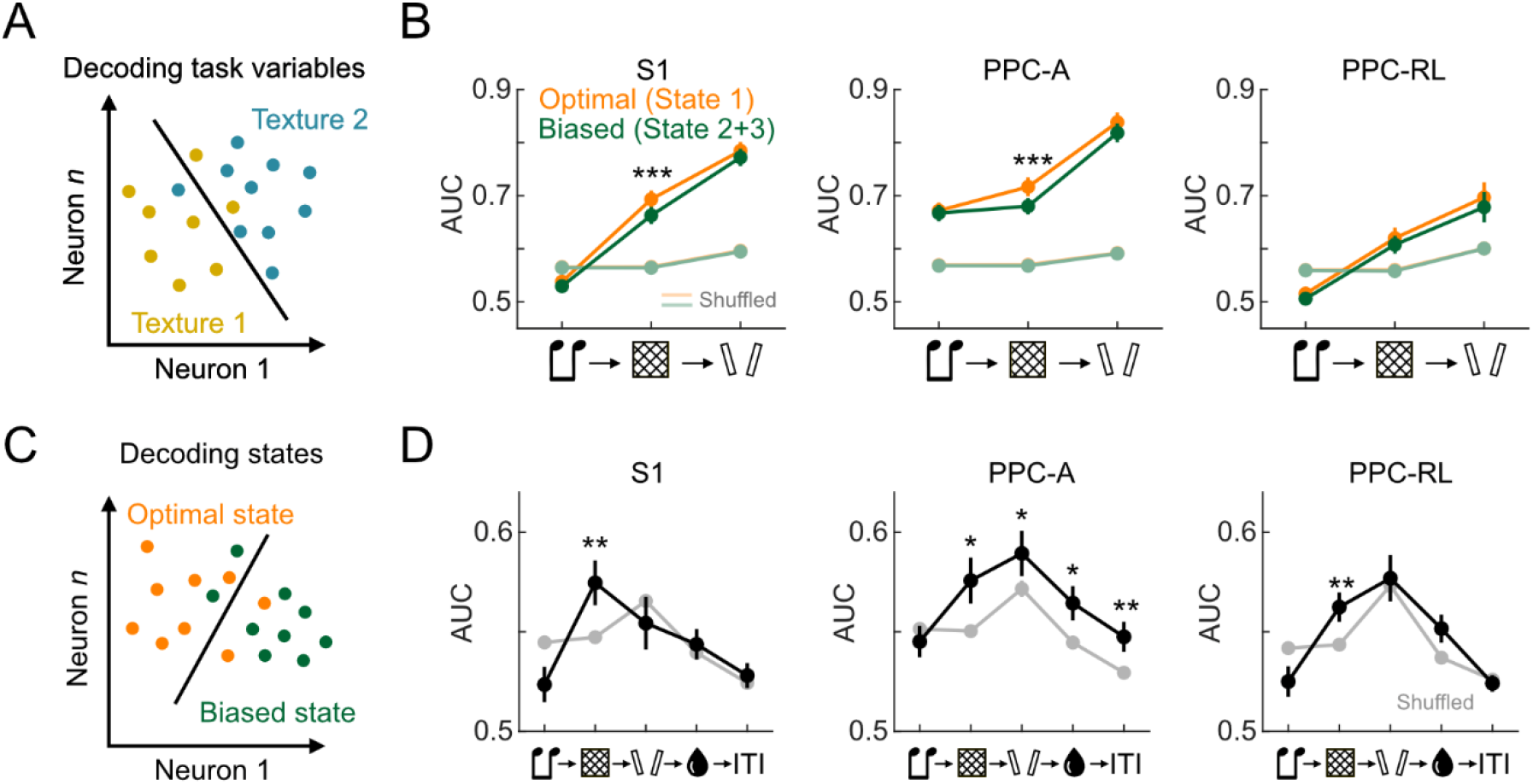
Task variable and behavioral state decoding from S1 and PPC activity. (**A**) Schematic of a linear decoder to decode task variables, e.g. texture stimulus. (**B**) Area-under-the-curve (AUC) for tone, texture and choice decoders across S1 and PPC areas. Decoding was performed using data from the corresponding window (e.g. the tone decoder was trained in the tone window). Orange lines represent decoding performance in state 1; green lines represent decoding performance in combined state 2 and 3 trials; light orange and light green lines represent the 95% quantile of shuffled data. (**C**) Schematic of a linear decoder to decode behavioral state. (**D**) State decoding AUC in separate task windows across S1 and PPC areas. Black lines represent decoder performance with real data; gray lines represent 95% quantile of shuffled data results. (S1: 13 mice, 53 sessions; PPC-A: 5 mice, 28 sessions; PPC-RL: 8 mice, 25 sessions; \**p*<0.05; \*\**p*<0.01; \*\*\**p*<0.001)

### State-dependent cross-area communication between S1 and PPC

During the task, S1 and PPC communicate continuously to support the transformation of sensory information into choice and action. Specifically, our previous studies showed that in this task PPC-A is more involved than PPC-RL in processing sensory information from S1^22,23^. We next asked whether S1-PPC interactions are different across behavioral states. To answer this question, we applied reduced-rank regression (RRR) to analyze the interactions between S1 and PPC, using population activity in one area to linearly predict population activity in the other area through a low-dimensional mapping^30,31^ (Figure 5A; Methods). In RRR, the regression coefficient matrix is constrained to be low rank, such that a small number of dimensions in the source population explains activity in the target population. These dimensions capture the components of population activity that best predict the target area and are ordered by their contribution to prediction performance. Importantly, RRR does not infer causal or synaptic interaction between brain areas; instead, it quantifies low-dimensional predictive relationships in population activity, which may reflect shared signals, common input, or inter-area communication. To reduce the contribution of common inputs that arise from shared task structure, and to improve computational stability, we first performed PCA on each population and kept the top 30 principal components (PCs), then subtracted condition averages from the population activity and performed RRR on the residual activity. This procedure is designed to retain inter-area communication patterns beyond the stimulus-driven mean response for each trial type. We repeated the analysis on bootstrapped sets of correct trials with matched trial numbers across states and averaged the results.

**Figure 5.**
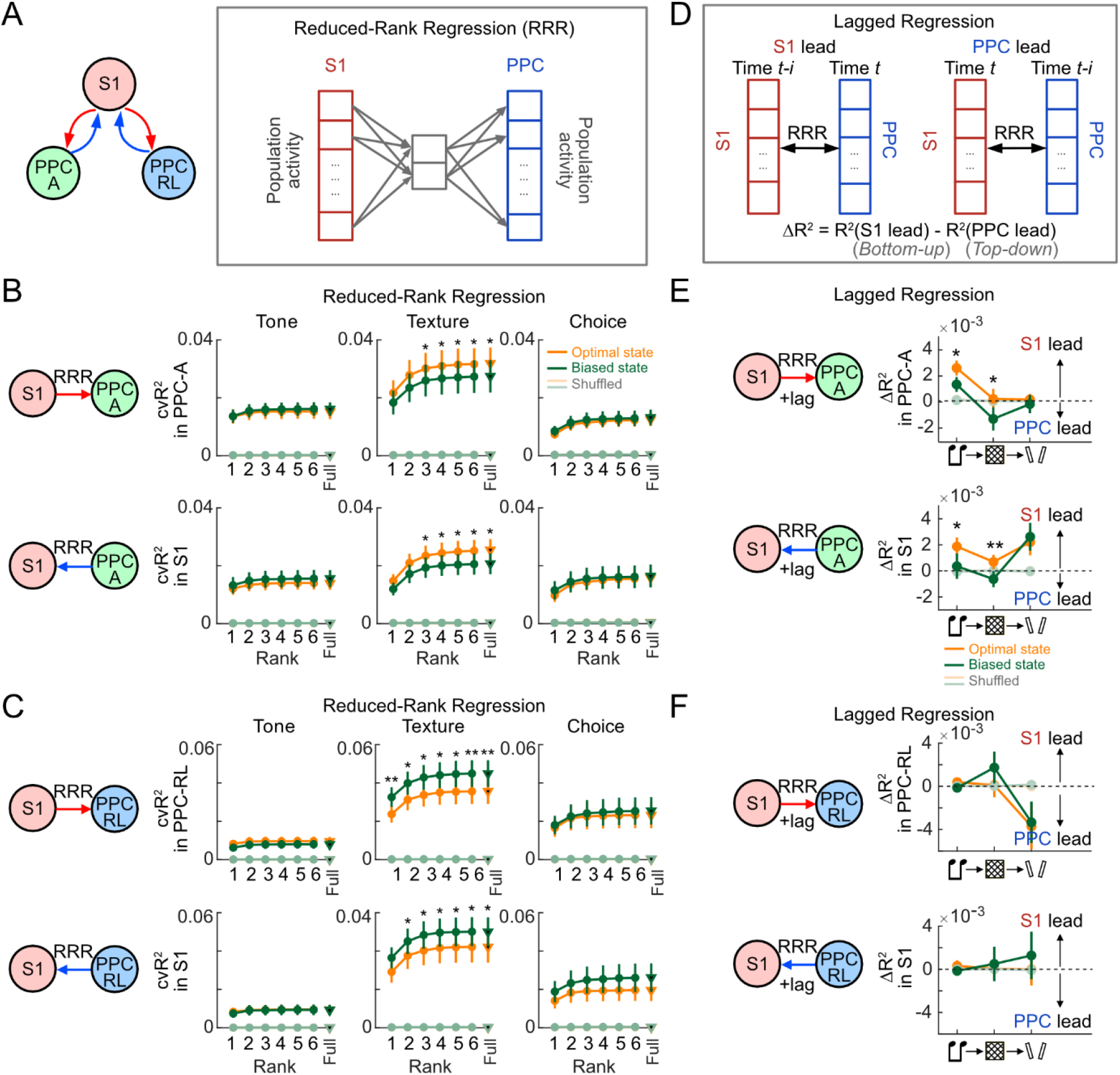
Reduced-rank regression analysis of S1-PPC communication across behavioral states. (**A**) Schematic of reduced-rank regression (RRR) analysis for S1 and PPC interactions, where we simultaneously imaged from S1 and PPC-A, or S1 and PPC-RL. RRR identifies a set of low-rank linear regression coefficients using the activity in S1 to predict the activity in area PPC, or using PPC activity to predict S1 activity. (**B**) RRR analysis between S1 and PPC-A. Top panels: cross-validated R^2^ (cvR^2^) from RRR models using S1 to predict PPC-A activity with varying model ranks. Triangles represent the full rank results. Light colored lines represent shuffled results. Bottom panels: cross-validated R^2^ from RRR models using PPC-A to predict S1 activity. RRR in both directions showed similar results. (**C**) RRR analysis between S1 and PPC-RL. Top panels: cross-validated R^2^ from RRR models using S1 to predict PPC-RL activity with varying model ranks. Bottom panels: cross-validated R^2^ from RRR models using PPC-RL to predict S1 activity. RRR in both directions showed similar results. (**D**) Schematic of RRR analysis with temporal lags. RRR was performed with an additional lag (up to ±0.5 second) in the source area. The R^2^ differences between S1-lead models and PPC-lead models were computed to represent the top-down and bottom-up influences. (**E**) Lagged RRR for S1 and PPC-A interactions, using S1 to predict PPC-A activity (top), or using PPC-A to predict S1 activity (bottom). ΔR^2^ was higher in the optimal performing state (orange lines) than the biased state (green lines), indicating stronger bottom-up influences. (**F**) Lagged RRR for S1 and PPC-RL interactions, using S1 to predict PPC-RL activity (top), or using PPC-RL to predict S1 activity (bottom). (S1 and PPC-A analyses: 5 mice, 28 sessions; S1 and PPC-RL analyses: 8 mice, 25 sessions; \**p*<0.05; \*\**p*<0.01)

We performed RRR using S1 to predict PPC activity, or PPC to predict S1 activity, while systematically varying the rank of the regression model (Figure 5B-C). Across task windows, the communication strength between S1 and both PPC subregions, measured by cross-validated R^2^ (cvR^2^), was highest during texture presentation. This communication resided in a low-dimensional subspace, as the regression performance already saturated at low ranks, reaching comparable performance to the full-rank model performance. Notably, communication between S1 and PPC-A was stronger during the optimal state than during the biased state, whereas communication between S1 and PPC-RL was weaker during the optimal state than during the biased state (Figure 5B-C). Canonical correlation analysis (CCA) recapitulated these trends in S1-PPC communication (Figure 5 – figure supplement 1). These results indicate state-dependent differences in S1-PPC communication for both PPC subregions and suggest preferential routing of task-relevant sensory information through PPC-A in the optimal-performing behavioral state.

We next asked whether communication between S1 and PPC showed different temporal patterns across behavioral states, consistent with either top-down information flow, with PPC activity preceding S1 activity, or bottom-up information flow, with S1 activity preceding PPC activity. To assess these temporal relationships, we introduced a temporal lag in the reduced-rank regression model by shifting activity in the source area. We then computed the difference in explained variance (ΔR^2^) between the S1-lead models, in which S1 activity preceded PPC activity, and PPC-lead models, in which PPC activity preceded S1 activity (Figure 5D). We performed this lagged RRR analysis in both directions, using S1 to predict PPC activity and PPC to predict S1 activity. During the optimal state, models in which S1 activity preceded PPC-A activity showed higher regression performance than models in which PPC-A activity preceded S1 activity, resulting in a positive ΔR^2^. This ΔR^2^ was larger compared to the biased state, suggesting stronger bottom-up information flow from S1 to PPC-A during optimal behavioral performance (Figure 5E). In contrast, we did not detect a significant temporal asymmetry between S1 and PPC-RL (Figure 5F). We verified these findings using CCA analysis and found a similar temporal pattern (Figure 5 – figure supplement 1). Together, these results suggest that optimal task performance is associated with stronger bottom-up information flow from S1 to PPC-A.

### Communication between task-related and task-orthogonal subspaces in S1 and PPC

During behavioral tasks, cortical areas not only carry task-relevant information, but also activity that relates to ongoing behavioral state, movement, and other task-independent factors. We therefore asked whether communication of task-related and task-orthogonal activity differed between the optimal and the biased state. To address this question, we defined a task-related subspace using the axes of linear decoders trained to discriminate tone, texture, and choice information. We then defined task-orthogonal subspace as the null space of the task-related subspace (Figure 6A; Methods). To separate task-related and task-orthogonal activity in each population, we projected the activity of S1 and PPC onto their task-related subspace axes to represent task-related activity; likewise, we projected activity onto the task-orthogonal space to represent activity orthogonal to those task dimensions (Figure 6B). In the task-orthogonal subspace, PPC-A activity still carried information about behavioral state, as linear decoders were able to read out behavioral state from PPC-A task-orthogonal activity, whereas S1 and PPC-RL did not carry decodable behavioral state information (Figure 6C). Thus, state-related signals in PPC-A were not confined to the decoder-defined task dimensions.

**Figure 6.**
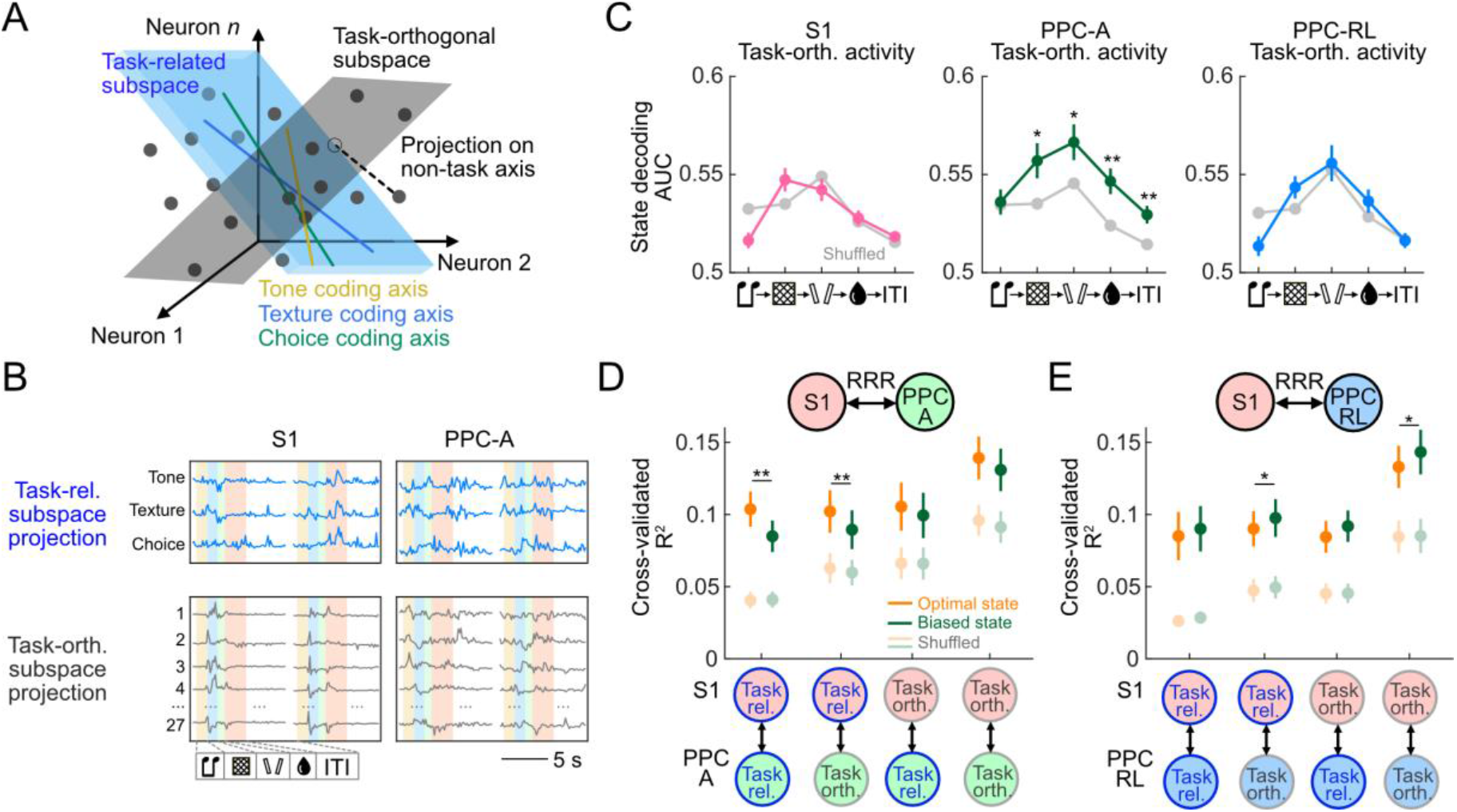
Communication between task-related and task-orthogonal activity in S1 and PPC. (**A**) Schematic of defining task subspace and task-orthogonal subspace. Task subspace was defined using the tone, texture, and choice coding axes; task-orthogonal subspace was defined as the null space of the task subspace. (**B**) Example of task-related (“Task-rel.”) subspace projection activity and task-orthogonal (“Task-orth.”) subspace projection activity in S1 and PPC-A. The task-related and task-orthogonal subspace were defined using top 30 PC activity. (**C**) Decoding performance for decoding the behavioral state using the task-orthogonal activity in S1, PPC-RL and PPC-A, across task windows. (**D**) Cross-validated R^2^ for RRR analyses between the task-related and task-orthogonal activity in S1 and PPC-A (left), or S1 and PPC-RL (right). The R^2^ was computed using texture window activity, and was the average of RRR models using S1 to predict PPC activity, and using PPC to predict S1 activity. (S1: 13 mice, 53 sessions; PPC-A: 5 mice, 28 sessions; PPC-RL: 8 mice, 25 sessions; S1 and PPC-A analyses: 5 mice, 28 sessions; S1 and PPC-RL analyses: 8 mice, 25 sessions; \**p*<0.05; \*\**p*<0.01; \*\*\**p*<0.001)

We next asked whether communication between S1 and PPC differed for task-related and task-orthogonal activity across behavioral states. To test this idea, we performed reduced-rank regression separately on task-related and task-orthogonal activity in each area. For S1 and PPC-A interactions, task-related activity in S1 during the texture window explained both task-related and task-orthogonal activity in PPC-A better in the optimal state than in the biased state, whereas task-orthogonal activity in S1 explained PPC-A task-related and task-orthogonal activity similarly across states (Figure 6D). This pattern suggests a stronger transmission of S1 task-related signals to PPC-A during the optimal state, indicating that task-related sensory signals in S1 may interact with state-related or other task-independent signals in PPC-A. In contrast, both task-related and task-orthogonal activity in S1 during the texture window explained task-orthogonal activity in PPC-RL better during the biased state compared to the optimal state (Figure 6E). This result suggests shared state-dependent fluctuations rather than selective communication of task-relevant sensory information to PPC-RL. Communication in other task windows did not differ across states (Figure 6 –figure supplement 1). Taken together, these results indicate that the optimal performing state is associated with stronger communication of task-related activity between S1 and PPC-A, whereas biased states are associated with stronger communication of task-orthogonal activity between S1 and in PPC-RL.

## Discussion

Sensory-guided behavior depends not only on accurate sensory representations within specific areas, but also on how task-relevant information is communicated across cortical networks^17,32–36^. Using a multisensory choice task with simultaneous two-photon imaging in S1 and PPC, together with analysis using GLM-HMM-based latent-state inference, we found that the optimal-performing behavioral state was associated with stronger sensory representations in local populations and stronger bottom-up communication from S1 to PPC-A. In contrast, the two biased states were associated with weaker local sensory coding and different patterns of inter-area communication, with relatively stronger involvement of PPC-RL than PPC-A. These state-dependent differences were not restricted to task-related dimensions of population activity but also extended to task-orthogonal activity in PPC. Together, these findings suggest that different behavioral states are associated with differences in both local sensory encoding and S1-PPC communication (Figure 7). More broadly, the results support the idea that behavioral state influences not only representations within sensory cortex, but also along which inter-area pathway task-relevant sensory information between sensory and association areas is preferentially routed.

**Figure 7.**
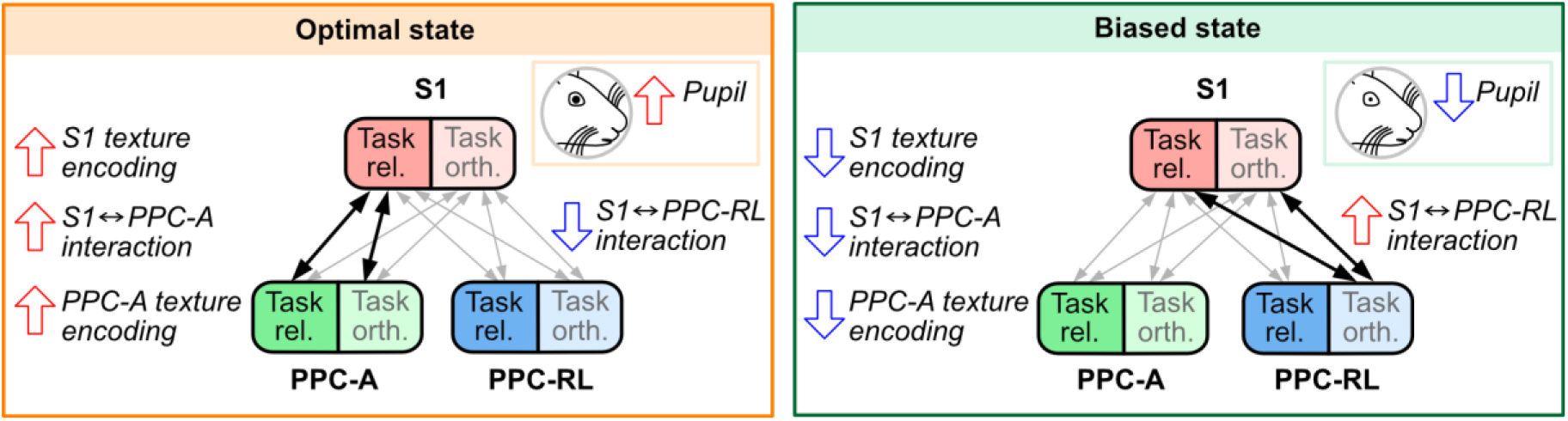
Summary diagram. A summary of findings in this study. Optimal state is associated with larger pupil size, stronger sensory representation in S1 and PPC-A, stronger bottom-up interaction from S1 to PPC-A, and stronger communication between S1 task activity and PPC-A activity. Biased state is associated with smaller pupil size, weaker sensory representation in S1 and PPC-A, stronger interaction between S1 and PPC-RL, and stronger communication between S1 and PPC-RL task-orthogonal activity.

Many studies have shown that PPC is involved in multiple steps of sensory processing and decision making. PPC receives visual, somatosensory, auditory, and egocentric inputs, and integrates information across sensory modalities and over time^37–44^. Beyond sensory information, PPC also carries signals related to choice, trial history, and movement, linking sensory inputs to decisions and actions^29,37,45,46^. Several studies suggest that PPC can maintain sensory information and relay it to frontal areas for decision-making^38,45^, particularly during the learning of tasks^47,48^. At the same time, the precise contribution of PPC appears to depend strongly on task demands. PPC is less critical in some simple sensory discriminations, but becomes more important when the task requires flexible sensorimotor mapping, memory, resolution of uncertain sensory evidence, or integration across time and modality^24,28,39,42,49^. Our previous work showed that in the present behavioral task, texture-period activity in both PPC-A and PPC-RL is required for optimal performance, and that both subregions remain necessary even when only tone or only texture cues are available^23^. The present results add behavioral state as another factor that shapes when and how PPC contributes to behavior.

A growing body of work shows that the rodent PPC contains functionally distinct subregions^1,37,50,51^. The anterior region PPC-A has been linked to visual and auditory processing, history-dependent signals, and movement-and posture-related activity^17,22,29,37^; the anteromedial region (area AM) encodes both vision and choice information and exhibits diverse coupling with posterior cortex^37,50^; the rostrolateral region PPC-RL stores visual and tactile information during the delay period and emerges as a key node during learning of sensorimotor tasks^17,32,40,47,48^. These observations support the idea that PPC is not a unitary structure, but a set of subregions whose contributions depend on task configuration, sensory modality, training history, and behavioral context^17,29,37,49^. Our results add behavioral state to this list. S1 communication with PPC-A and PPC-RL was relatively stronger for PPC-A during the optimal state and for PPC-RL during the biased states. Together with our previous finding that PPC-A communicates cross-modal sensory predictions to S1 in this task, the present results suggest that task configuration, learning, and behavioral state can all influence which S1-to-PPC pathway is preferentially recruited. This view suggests that functional specialization within PPC may contribute to functional pathways and flexible information routing across cortical networks.

Behavioral state is represented broadly across the brain, and can be multiplexed in separate channels independent of sensory representation^2,51–53^. Several mechanisms could contribute to state-dependent information processing. One possibility is selective information routing, in which sensory information is preferentially communicated through different cortical pathways depending on behavioral state^18^. Another possibility is that state changes alter the shared variability or synchrony across areas, thereby changing how effectively one population can influence another^24,54,55^. These two possibilities are not mutually exclusive. In our data, the analyses for task-related and task-orthogonal activity suggest that behavioral state affects both the population dynamics in PPC and the communication of task-relevant sensory signals to PPC. This implies that state dependence may not act only on sensory feature encoding, but also on the multi-regional population dynamics in which those signals are embedded. Many biological mechanisms could support these effects, including changes in arousal, attention, locomotion, and neuromodulation, all of which can alter gain and synchrony^56,57^. Neuromodulatory systems may be particularly important: for example, acetylcholine can enhance feedforward sensory input relative to feedback signals^58^, thus improving bottom-up transmission of sensory information, while noradrenergic signaling has been linked to rapid cortical state reconfiguration and large-scale network coordination^59,60^. Such mechanisms could allow behavioral state to shape both local sensory representations and inter-area communication on behavioral timescale. In addition, projection patterns between areas could support communication between areas so that relevant neuronal populations can work together to generate consistent neural code^24,61^. Our reduced-rank regression analyses do not distinguish whether the observed changes arise from direct inter-areal interactions, shared upstream drive, or parallel modulation of both. Future experiments that combine simultaneous recording with targeted perturbation of PPC subregions or state-regulating systems will be needed to separate these possibilities.

Behavioral state is multidimensional, including arousal, attention, motivation, movement, and strategy, and these factors can coexist at the same time^4,6^. As in previous studies^10,19,24,27^, we identified three behavioral states, one associated with accurate, stimulus-guided performance and two associated with an intrinsic choice bias to lick either right or left. The three latent states identified here should not be interpreted as pure or exhaustive state categories. Instead, they can be viewed as a simple set of discrete behavioral states inferred from choice patterns. Because these states were inferred from behavior rather than measured directly from neural or physiological variables, they likely compress multiple internal processes into a small number of discrete states. Previous work using a sound localization task showed that latent behavioral states were reflected more strongly in PPC than in auditory cortex^24^. Our findings are generally consistent with this view but also extend it in two ways. First, S1 was not state-invariant: sensory representations in S1 were weaker in the biased states than in the optimal state. Second, state-dependent differences were stronger in PPC-A and in S1-PPC communication than in S1 alone. These observations suggest that state-dependent behavior is not explained solely by local changes in sensory coding. Instead, behavioral state may act most strongly by regulating inter-area communication, e.g., whether sensory signals in primary cortex gain effective bottom-up access to neural circuits in association areas. This framework suggests that flexible behavior depends not simply on global changes in sensory gain, but also on selective recruitment of distinct inter-area pathways, which both shape and reflect the current behavioral state.

## Materials and methods

### Mice and dataset

In this study, we performed novel analysis on a previously published dataset from our 2025 study^23^. A summary of the animal and experimental procedure is provided below, and the full method can be found in the original study. Briefly, a total of 16 mice were included in this study. Both sexes were included in this study (8 male mice, 8 female mice). Mice belonged to one of the following transgenic strains: RasGRF2a-dCre;CamK2a-tTA;TITL-GCaMP6f (10 mice), GP5.17(C57BL/6J-Tg(Thy1-GCaMP6f)GP5.17Dkim/J, Jackson Laboratory 025393) (2 mice), Snap25-IRES2-Cre-D;CamK2a-tTA;TITL-GCaMP6f (1 mouse), RasGRF2a-dCre;tTA2-GCaMP6f (3 mice). All mice expressed GCaMP6f in layer 2/3 pyramidal neurons of the neocortex. One mouse was only used for behavioral modeling due to the insufficient cranial window quality for two-photon imaging. Imaging data was collected on 15 mice.

### Behavioral task

Behavioral training was conducted using custom-written LabVIEW software that controlled National Instruments devices. In the task, mice were head-fixed in a body-restraining tube and received auditory cues from bilateral speakers and tactile stimuli delivered to their whisker pad with sandpaper. Each trial started with a 0.5-second baseline period. Then, a pure tone (10 kHz or 18 kHz, six repetitions, 50-ms duration and 50-ms intervals) was played through the loudspeakers. One second after the tone onset, a sandpaper (P100 versus P1200, or P280 versus P800) mounted on a rotary motor carried by a linear motor stage moved into position, and the rotary motor swept the sandpaper (texture) onto the whisker pad of the mice. The sandpaper was presented for 1 second, after which a choice tone (two beeps at 3 kHz of 50-ms duration with 50-ms interval) was played to indicate the start of the choice window. Mice were free to lick the left or the right lick port during the following 2-s window, and the first lick during this choice window triggered the start of the reward window. If the mice made the correct choice, a drop of sugar water reward was delivered through the corresponding lick port. Incorrect choices were not punished. The intertrial interval was randomly distributed between 4 and 8 s.

During task training, tone-texture pairing was fixed. The low-frequency tone (10 kHz) was always followed by the rough texture (P100 or P280), whereas the high-frequency tone (18 kHz) was always followed by the smooth texture (P800 or P1200). Once mice became stable experts in this task (expert performance for at least three days), we introduced a small fraction (10–30%) of tone-texture mismatch trials, in which the tone was paired with the non-matching texture. In our previous study, we used the mismatch datasets to study predictive processing, but we did not include them in the present study for two reasons. First, the small number of mismatch trials did not provide sufficient data for GLM-HMM analysis. Second, we could not apply the same GLM-HMM framework to these trials because their sensory inputs were incongruent as the “stimulus” task variables. Accordingly, the dataset used in this study comprised sessions with only paired stimuli collected before mismatch trials were introduced.

We performed all training in the dark and continuously monitored mice with a CMOS infrared-sensitive camera (Basler acA1440–220um) under 850-nm infrared LED background illumination. A small UV LED (385 nm, Thorlabs LED385L) positioned close to the contralateral eye was used to constrain the pupil. We tracked pupil diameter by binarizing the pupil image and fitting an ellipse to the pupil region. Using manually drawn regions of interest (ROIs), we quantified body and face movement from changes in frame-to-frame correlation.

### Localizing S1 and PPC in the cranial window

To identify the precise locations of S1, PPC-A, and PPC-RL, we performed widefield sensory mapping and retinotopic mapping through the 5-mm cranial window. For sensory mapping, lightly anesthetized mice received contralateral visual, whisker, and hindlimb stimulation while widefield cortical activity was recorded. For retinotopic mapping, we presented drifting checkerboard stimuli moving in the four cardinal directions across the visual field. The sensory and retinotopic maps were aligned to the Allen Mouse Common Coordinate Framework to define the final recording sites within the cranial window.

### Two-area two-photon imaging

The imaging dataset was published in our previous study^23^. Briefly, imaging was conducted using a custom-built two-area two-photon microscope^62^. In each session, we simultaneously recorded population activity from layer 2/3 neurons in S1 and either PPC-A or PPC-RL. Typically, imaging covered a ∼450×500 µm field-of-view at 3 depths within L2/3 at 9.3 vol/s, yielding in ∼50 to ∼600 neurons per area. In the full dataset, 15 mice were imaged: 5 mice with simultaneous S1 and PPC-A recordings, and 10 mice with simultaneous S1 and PPC-RL recordings. For subsequent state-dependent analyses, we excluded sessions with low trial counts due to their low computational stability, and only analyzed sessions with a minimum of 15 trials from each state (state 1, or state 2 and 3 combined). After applying this criterion, the final dataset analyzed included 28 sessions for S1 and PPC-A from 5 mice, and 25 sessions for S1 and PPC-RL from 8 mice. Calcium traces were deconvolved using Suite2p^63^, and the deconvolved spike rates were used for all following analyses.

### GLM-HMM model selection

To identify behavioral states, we applied a hidden Markov model with Bernoulli Generalized Linear Model observations (GLM-HMM)^19,25^. For a K-state GLM-HMM, the model consists of a *K* × *K* state transition matrix *A*, and a set of GLM weights *w_k_* that map the input covariates to the choice probability for each state. In our study, we defined the input covariates as combinations of the following task variables: stimulus, previous trial choice, strategy (win-stay-lose-switch), previous trial reward, additional choice history from three preceding trials (“Prev. N choice”), additional reward history from three preceding trials (“Prev. N reward”), and a constant term (“Bias”). All task variables were represented as –1 or +1. The task variables were encoded into an *M* × *T* matrix, where *M* is the dimension of the task variables, and *T* is the total number of trials.

To select the most relevant task features for predicting behavioral choice, and to identify the best number of states, we first fit the GLM-HMM on all sessions from all mice. During task sessions, mice did not lick in a small fraction of trials (“miss trials”); in these trials, choice could not be represented as a binary variable, where –1 indicates left choice and 1 indicates right choice. To account for the non-binary choices in these miss trials, we fitted a multinomial GLM-HMM. However, the number of miss trials was too low to support stable model fitting. We therefore excluded all miss trials and used a Bernoulli GLM-HMM. We fit models using the expectation-maximization (EM) algorithm and inferred posterior state probabilities with the forward-backward algorithm. To avoid carryover across sessions or long gaps in valid choices, we reinitialized the posterior state probabilities at the start of each session and after blocks of at least five consecutive miss trials. For model fitting, we initialized each row of the transition matrix with a Dirichlet distribution that favored self-transitions (diagonal concentration = 5, off-diagonal concentration = 1). All model fitting was performed with 10-fold cross validation, where folds were split within each session in contiguous blocks. Withholding 10% of sessions instead of 10% trials within each session did not affect model performance and identified states. Model hyperparameters were selected with a grid search, where we set the number of states to be [1, 2, 3, 4, 5, 6], and the L2 regularization parameter lambda to be [0.1, 0.5, 1.0, 2.0, 5.0, 10.0]. As a baseline, we additionally fit a 1-state bias-only Bernoulli GLM. Model performance was evaluated using log-likelihood (LL) expressed in bit per trial (bpt), defined as following^19,27^:

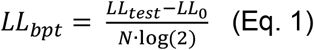

Here, *N* is the number of trials in the test set, *LL_test_* is the log-likelihood of the test set, and *LL*_0_ is the log-likelihood of the same test set using bias-only Bernoulli GLM. Because the EM algorithm does not guarantee convergence to the global maximum likelihood, we ran EM 20 times for each fold and selected the model with the highest model fitting performance. We confirmed that the top models all converged to the same solution. Additionally, we also computed the choice prediction accuracy of the models by comparing the model prediction of choice with the behavioral choice data. With these steps, we selected the model input parameters to be the stimulus and bias terms, and that optimal number of states to be three. We refer to the transition matrix and the GLM weights from all mice and sessions as *A_global_* ^and^ *^w^global*.

To refine the model and better capture behavioral state across learning, we considered two approaches. The first approach was to train mouse-specific models initialized by *A_global_* and *w_global_*. This approach assumes that each individual mouse has its own set of latent states, reflected by mouse-specific GLM weights, and that the transitions between these states are invariant across learning phases. The second approach was to train learning-phase-specific models pooled across mice using fixed *w_global_*, but learning-specific transition matrices. This approach assumes that mice share common behavioral states, but the occupancies and transitions of these states are different across learning phases. As previous work has shown that mouse decision-making can be described by a small but fixed set of latent strategies shared across animals, we chose to keep the GLM weights fixed across mice and fit learning-phase-specific transition matrices so that states remained comparable across mice and sessions^27^. With this approach, the states were directly comparable across mice and sessions. As in our previous study and other studies using similar paradigms^4,23^, we defined three learning phases: naïve (session average performance <55%), learning (performance between 55% and 75%), and expert (performance ≥ 75%). Model training was performed as above with 5-fold cross validation. This model tuning yielded model prediction accuracy of 62.5%, 64.7% and 78.0% for the naïve, learning, and expert phases, respectively, indicating above-chance prediction in all phases and improved predictability with learning.

Within each state, mice made both correct and incorrect choices. To reduce the confounds of different stimulus-choice contingencies and trial outcomes, we only used the correct trials for all subsequent analysis of the neural data. We combined state 2 and 3 as a single biased state and sampled equal numbers of trials for the optimal performing state (state 1) and the biased state. This sampling procedure was repeated 20 times to minimize the bias from trial selection. To ensure computational stability, sessions with fewer than 15 trials per state were excluded from further analyses. We only used data from learning and expert phases because state 1 in naïve mice likely reflect different underlying processing, as these mice have not yet learnt the behavioral task.

### Decoding analyses for task variables and behavioral states

To decode task variables from population activity, linear support vector machines (SVMs) were used to decode three task variables: tone, texture, and choice. Each decoder was trained using population activity during the corresponding task window, that is, tone decoding using activity during the tone window, texture decoding during the text window, and choice decoding during the choice window. Each decoder was trained with two class labels, for example, tone 1 versus tone 2, using pooled frames across trials. As each neuronal population can contain up to 700 neurons, whereas the numbers of trials and time points were limited, we first performed a dimensionality reduction on the population activity using principal component analysis (PCA), and kept the top 30 PCs for each population as in our previous studies^22,23^. Only correct trials were used for decoding. To avoid overfitting, we regularized the SVM coefficients with an L2 penalty. The regularization term was cross-validated in a log space over seven values ranging from 10^−5^ to 10^1^. To avoid overfitting due to unbalanced class number, a misclassification cost inversely proportional to the number of trials in each class was also used. All decoders were trained using 5-fold cross validation.

Shuffled decoders were trained by shuffling the class labels using the same procedure as above. The shuffling procedure was repeated 100 times. To evaluate the decoder performance, the receiver operating characteristic curve and the area under the curve (AUC) were calculated using standard approaches, and shuffled AUC threshold was defined as the 95% quantile of shuffled distribution.

To decode behavioral state, we sampled equal number of trials from the optimal and the biased states, and trained decoders to classify the state labels using population activity from separate task windows. This trial sampling was repeated 20 times, and the results were averaged across repeats. Decoder and shuffled decoder training were performed using the same 5-fold cross validation as described above.

### Reduced-rank regression

We used reduced-rank regression (RRR) to quantify cross-area population coupling between S1 and PPC, following previous work^31^. As in the decoding analysis, the top 30 PCs from each population and matching numbers of correct trials (resampled 20 times, as described above) were used for analysis. To minimize the impact of shared task-related activity and to capture trial-to-trial fluctuations, the trial-type-averaged population activity was subtracted from each trial, and the residual activity was used for all analyses. RRR models were then fitted while varying the rank of the regression coefficient matrix. To reduce overfitting, we applied ridge regression and cross-validated the penalization parameter λ. Using the selected λ, full-rank ridge regression models (“full model”) were trained, and reduced-rank models were computed by constraining the coefficient matrix to ranks1-6. A rank-0 model was used as a null model, corresponding to the absence of cross-area predictive structure.

For each reduced-rank model, performance was quantified by subtracting the cross-validated R^2^ of the rank-0 model from that of the model at each rank. For shuffled data, we randomized the trial correspondence between the two areas by permuting trial order in one area 100 times while preserving the within-area activity structure and trial-type composition, and computed the cross-validated R^2^. RRR models were trained in both directions, using S1 activity to predict PPC-A or PPC-RL activity, and using PPC activity to predict S1 activity.

For lagged RRR, the activity of the source population (regressor) was shifted from –5 frames to +5 frames relative to the target population, corresponding to ±0.5 s. Negative lags indicated that source activity preceded the target activity, whereas positive lags indicated that source activity followed target activity. At each lag, we chose rank=5 to compute RRR models, as the regression performance plateaued at rank 5, reaching a comparable level as the full-rank model performance (see Figure 5BC). Cross-validated R^2^ was computed as above, and R^2^ values were averaged over lags up to 0.2 s for each lead direction. The final directional asymmetry metric was defined as ΔR^2^ = R^2^(S1 lead) – R^2^(PPC lead).

### Canonical correlation analysis

Canonical correlation analysis (CCA) was performed as an alternative measure of cross-area coupling between S1 and PPC. As CCA is more prone to overfitting, we used the top 10 principal components from each population. Same as the RRR analyses, we sampled matched numbers of correct trials from each state, and subtracted trial-type-averaged activity from each trial. CCA was applied to the residual population activity to identify pairs of population dimensions with maximal correlation across areas. The strength of cross-area coupling was quantified as the canonical correlation of the first canonical dimension in the held-out test set using 5-fold cross-validation. Shuffled controls were generated by permuting trial correspondence between the two areas while preserving within-area activity structure.

### Task-related and task-orthogonal activity

Task-related and task-orthogonal population activity were separated using the weight vectors of the linear decoders for tone, texture, and choice. For each recorded population, these three decoder weight vectors were computed as above in the 30-PC space using correct trials, and their span defined a task-related subspace. Single-trial population activity was projected onto this subspace to obtain task-related activity. A complementary task-orthogonal subspace was then defined as the null space of the matrix formed by the three decoder weight vectors, and activity projected onto this null space was defined as task-orthogonal activity. All projections were performed separately for each recorded population.

The decoder-defined task-related and task-orthogonal subspaces were constructed using all correct trials, and subsequent decoding and reduced-rank regression analyses on the projected activity were then performed using the same sampling and cross-validation procedures described above.

### Quantification and statistical analysis

All statistical analysis was done in MATLAB. In general, Wilcoxon signed-rank test was used for paired samples, and Wilcoxon Rank Sum test was used for non-paired samples. No normality test was performed since these tests do not assume normality. Two-sided tests were performed unless otherwise indicated. No statistical methods were used to pre– determine sample sizes. Error bars represent mean±SEM. P-values were corrected for false discovery rate (FDR) < 0.05 for multiple testing, using MATLAB function mafdr. Mice were randomly assigned to imaging groups (S1 and PPC-A or S1 and PPC-RL imaging). Stimulus presentation during imaging was fully randomized. Data collection and analysis were not performed blind to the conditions of the experiments.

## Acknowledgement

We thank Philipp Bethge for managing the transgenic mouse lines, Fabian Voigt and Hansjörg Kasper for help with optics, and Martin Wieckhorst for the behavior training software. We thank Alex Kwan, Luca Godenzini, and Christopher Lewis for their comments on the manuscript.

## Additional Information

### Funding

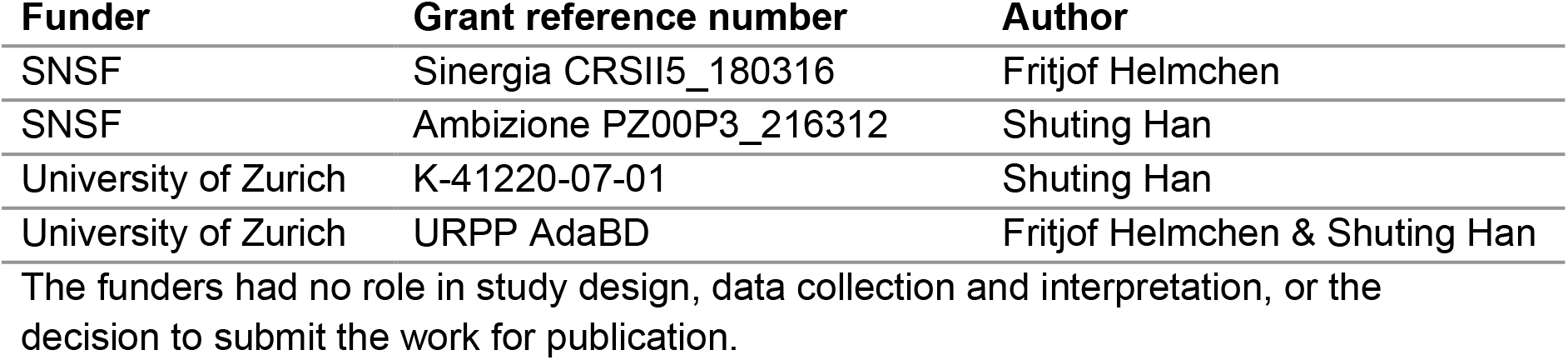

### Author Contribution

S.H. and F.H. conceived the study and designed the experiments. S.H. performed the experiments and analyzed the data. S.H. and F.H. wrote the manuscript.

## Additional Files

### Code availability

Data processing and analysis code is available at: https://github.com/HelmchenLabSoftware/multiarea_behavior_state_manuscript.

## Figure Supplements

**Figure 2 – figure Supplement 1.**
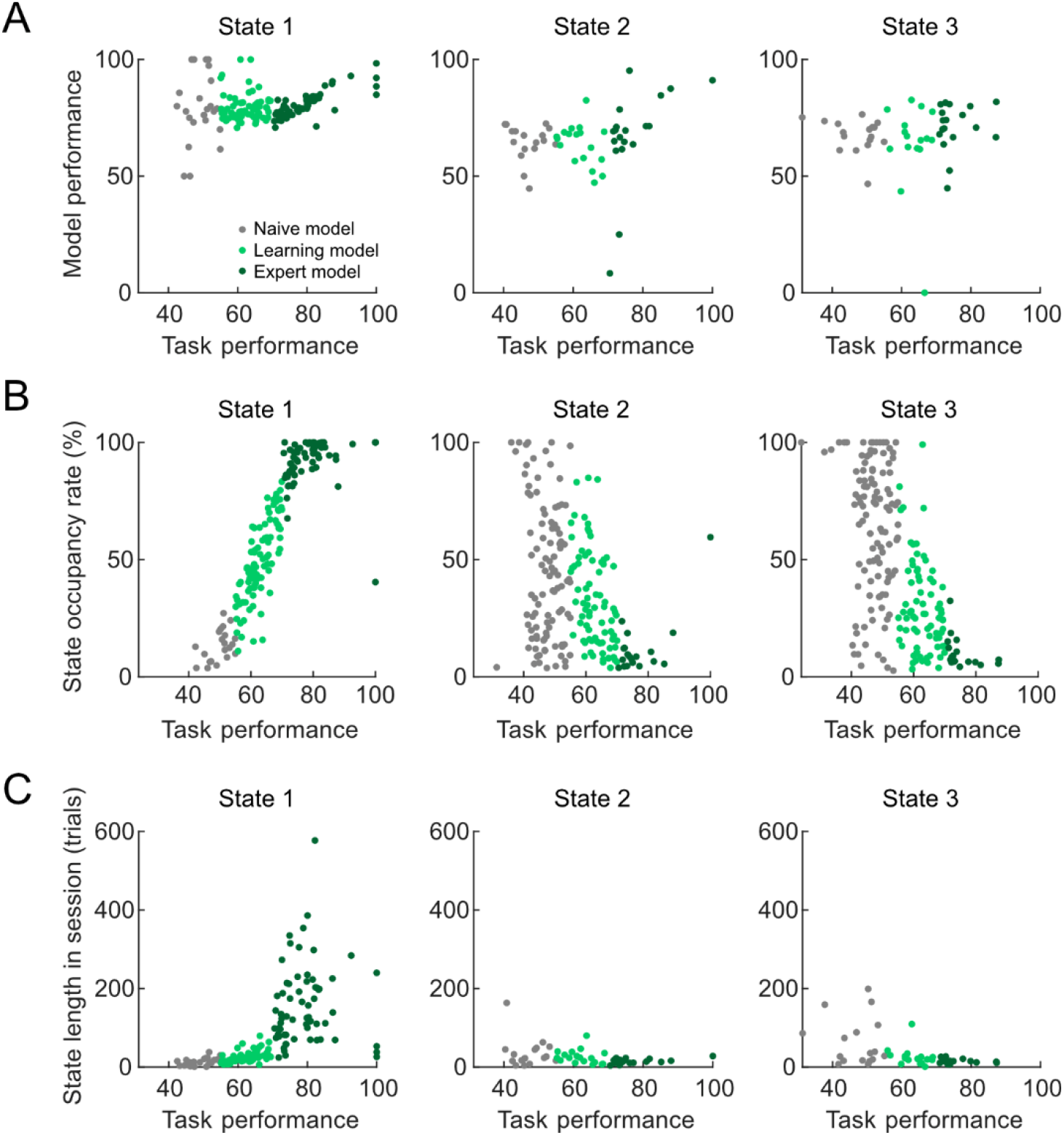
State characterization across learning phases. (**A**) Relationship between the task performance of mice and GLM-HMM prediction performance. Each dot represents a session; the color of the dots represents the learning phase. State 1 is the optimal state; state 2 and 3 are the biased states. (**B**) Relationship between the task performance of mice and the state occupancy rate within sessions. State occupancy rate was computed as the percentage of trials spent in each state. As mice learnt the task, state 1 occupancy rate increased, whereas state 2 and 3 occupancy rate dropped. (**C**) Relationship between the task performance of mice and the state length in each session. State length was computed as the number of continuous trials belonging to each state. As mice learnt the task, state 1 length increased, whereas state 2 and 3 lengths slightly dropped. (n=16 mice, 278 sessions; naïve: 117 sessions, learning: 96 sessions, expert: 65 sessions)

**Figure 4 – figure supplement 1.**
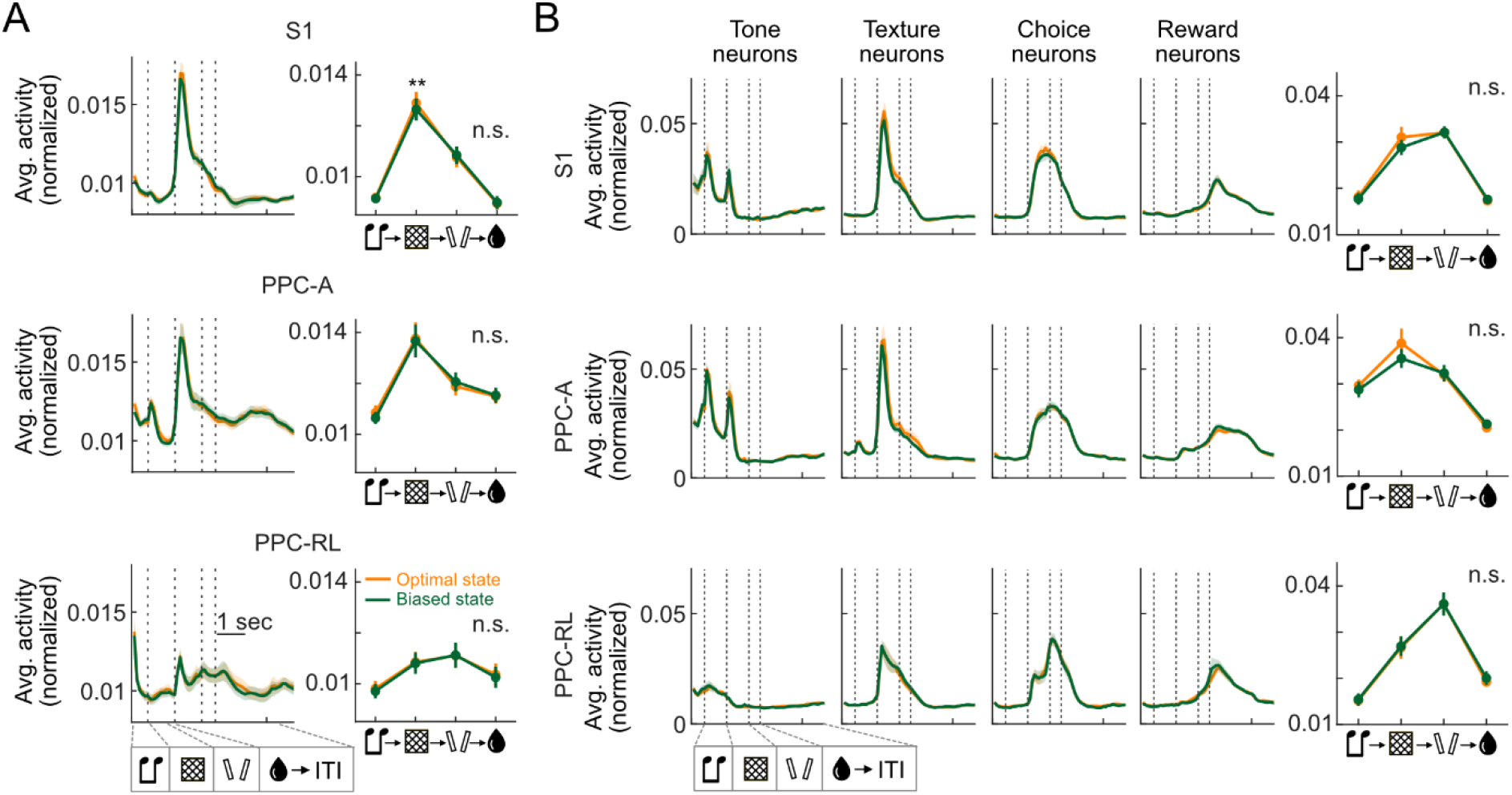
Population activity average in S1 and PPC across behavioral states. (**A**) Left: population activity average in S1, PPC-A, and PPC-RL over the task epochs. Right: average activity in each task window across behavioral states. (**B**) Left panels: average activity of task-tuned neurons for each task epoch and each area. Task-tuned neurons were identified by comparing each neuron’s activity within the specified task window with activity at randomly sampled time points outside that window, using matched sample numbers. Neurons with significantly higher activity within a task window than outside of the task window were identified as task neurons for that window. Right: average activity for task-tuned neurons in their corresponding task window. (S1: 13 mice, 53 sessions; PPC-A: 5 mice, 28 sessions; PPC-RL: 8 mice, 25 sessions; \*\**p*<0.01; n.s.: not significant).

**Figure 5 – figure supplement 1.**
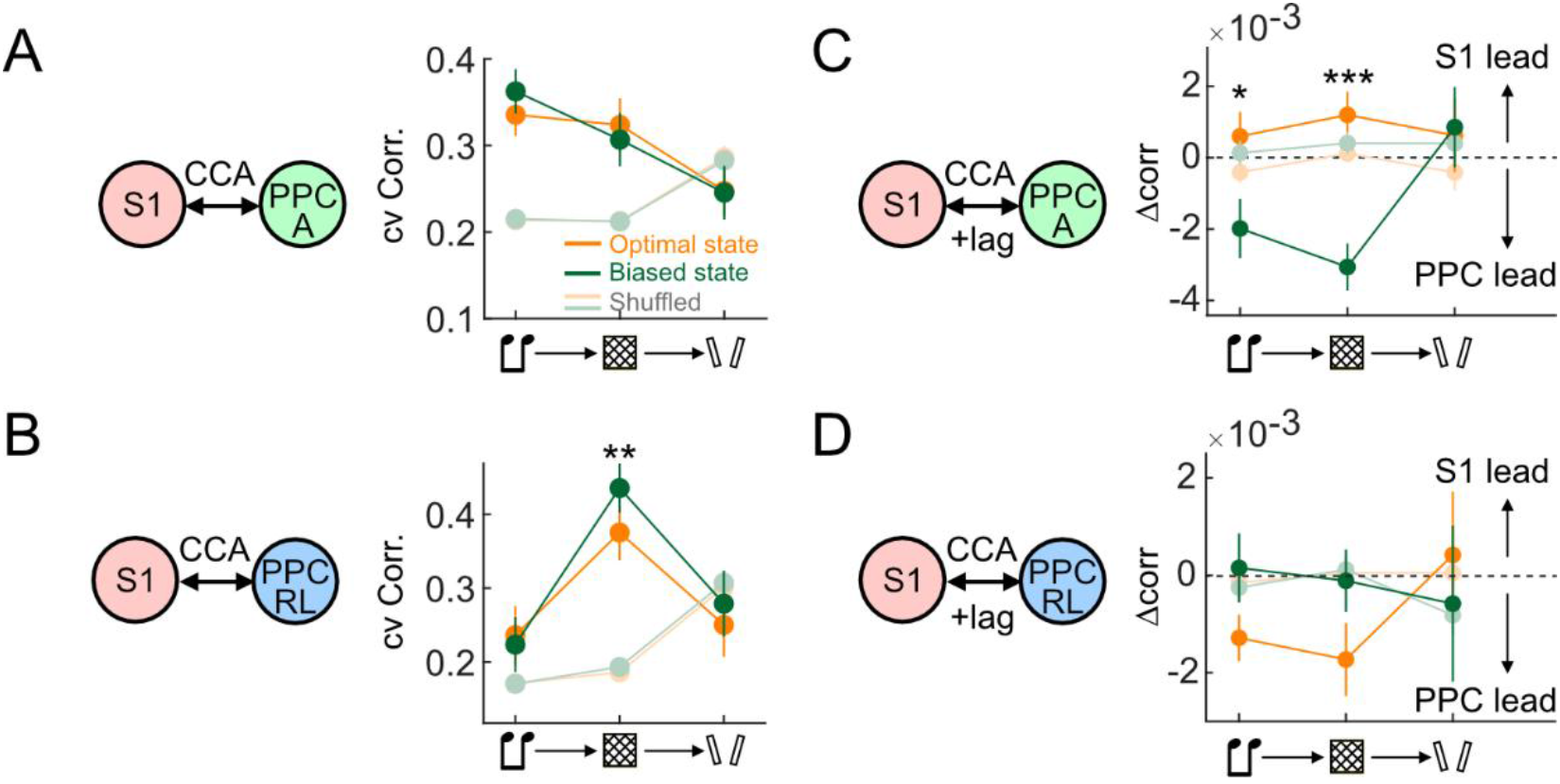
CCA analysis of S1-PPC communication across behavioral states. (**A**) Cross-validated CCA correlation between S1 and PPC-A across task windows and behavioral states. (**B**) Cross-validated CCA correlation between S1 and PPC-RL across task windows and behavioral states. (**C**) CCA analysis with temporal lags for S1 and PPC-A interactions. Similar to the RRR analysis, the top CCA correlation differences between S1-lead models and PPC-lead models were computed to represent the top-down and bottom-up influences. This Δcorr was higher in optimal performing state (orange lines) than the biased state (green lines), indicating stronger bottom-up influences. Note that unlike RRR, CCA is symmetric and treats S1 and PPC populations equally, therefore the directionality was computed using a single lagged CCA analysis instead of separate S1-to-PPC and PPC-to-S1 regression models. (**D**) Same as in C, but for S1 and PPC-RL interactions. (S1 and PPC-A analyses: 5 mice, 28 sessions; S1 and PPC-RL analyses: 8 mice, 25 sessions; \**p*<0.05; \*\**p*<0.01; \*\*\**p*<0.001)

**Figure 6 – figure supplement 1.**
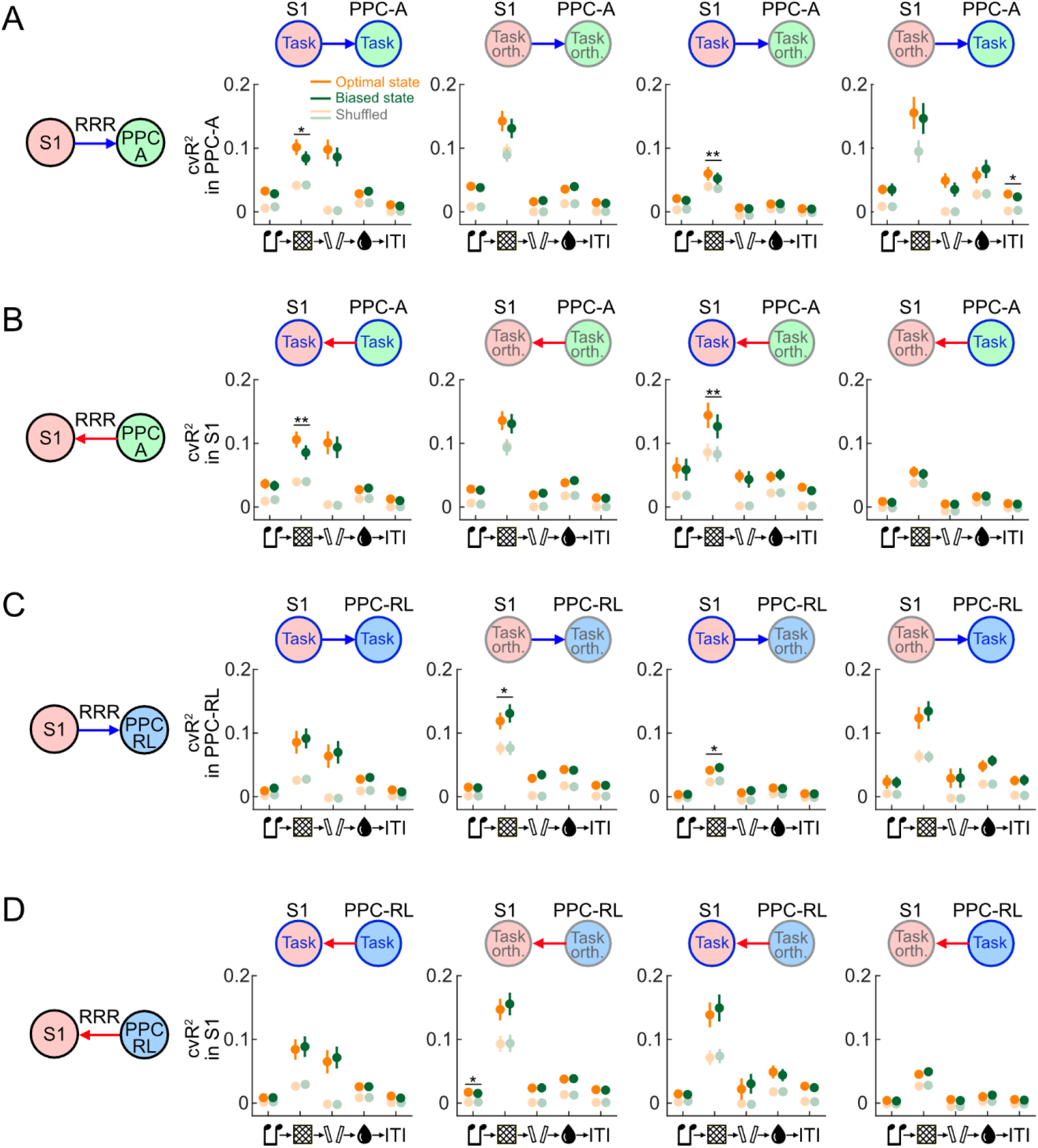
Bi-directional RRR between task-related and task-orthogonal activity in S1 and PPC. (**A**) Cross-validated R^2^ (cvR^2^) for RRR analyses using S1 to predict PPC-A activity for task-related and task-orthogonal activity, across all task windows. (**B**) Same as in A but using PPC-A to predict S1 activity. (**C**) Cross-validated R^2^ for RRR analyses using S1 to predict PPC-RL activity for task-related and task-orthogonal activity, across all task windows. (**D**) Same as in C but using PPC-RL to predict S1 activity.

## References

1. Biswas, R., Senevirathne, H.W., Wang, Y., Zhang, J., Mukherjee, S., and Abbasi-Asl, R. (2026). State-Dependent Organization of Microscale Functional Circuitry in Visual Cortex. bioRxiv. 10.64898/2026.03.31.715474.

2. Gardères, P.-M., Le Gal, S., Rousseau, C., Mamane, A., Ganea, D.A., and Haiss, F. (2024). Coexistence of state, choice, and sensory integration coding in barrel cortex LII/III. Nat. Commun. 15, 4782. 10.1038/s41467-024-49129-9.

3. Hulsey, D., Zumwalt, K., Mazzucato, L., McCormick, D.A., and Jaramillo, S. (2024). Decision-making dynamics are predicted by arousal and uninstructed movements. Cell Rep. 43, 113709. 10.1016/J.CELREP.2024.113709.

4. Abdolrahmani, M., Lyamzin, D.R., Aoki, R., and Benucci, A. (2021). Attention separates sensory and motor signals in the mouse visual cortex. Cell Rep. 36, 109377. 10.1016/j.celrep.2021.109377.

5. Ruff, D.A., and Cohen, M.R. (2019). Simultaneous multi-area recordings suggest that attention improves performance by reshaping stimulus representations. Nat. Neurosci. 22, 1669–1676. 10.1038/s41593-019-0477-1.

6. Clancy, K.B., Orsolic, I., and Mrsic-Flogel, T.D. (2019). Locomotion-dependent remapping of distributed cortical networks. Nat. Neurosci. 22, 778–786. 10.1038/s41593-019-0357-8.

7. Niell, C.M., and Stryker, M.P. (2010). Modulation of Visual Responses by Behavioral State in Mouse Visual Cortex. Neuron 65, 472–479. 10.1016/j.neuron.2010.01.033.

8. Morandell, K., Yin, A., Rio, R.T. Del, and Schneider, D.M. (2024). Movement-Related Modulation in Mouse Auditory Cortex Is Widespread Yet Locally Diverse. J. Neurosci. 44. 10.1523/JNEUROSCI.1227-23.2024.

9. Matteucci, G., Guyoton, M., Mayrhofer, J.M., Auffret, M., Foustoukos, G., Petersen, C.C.H., and El-Boustani, S. (2022). Cortical sensory processing across motivational states during goal-directed behavior. Neuron 110, 4176–4193.e10. 10.1016/j.neuron.2022.09.032.

10. Yin, C., Melin, M.D., Rojas-Bowe, G., Sun, X.R., Couto, J., Gluf, S., Kostiuk, A., Musall, S., and Churchland, A.K. (2025). Spontaneous movements and their relationship to neural activity fluctuate with latent engagement states. Neuron 113, 3048–3063.e5. 10.1016/j.neuron.2025.06.001.

11. Russell, L.E., Fişek, M., Yang, Z., Tan, L.P., Packer, A.M., Dalgleish, H.W.P., Chettih, S.N., Harvey, C.D., and Häusser, M. (2024). The influence of cortical activity on perception depends on behavioral state and sensory context. Nat. Commun. 15, 2456. 10.1038/s41467-024-46484-5.

12. McGinley, M.J., David, S. V., and McCormick, D.A. (2015). Cortical Membrane Potential Signature of Optimal States for Sensory Signal Detection. Neuron 87, 179–192. 10.1016/J.NEURON.2015.05.038.

13. McGinley, M.J., Vinck, M., Reimer, J., Batista-Brito, R., Zagha, E., Cadwell, C.R., Tolias, A.S., Cardin, J.A., and McCormick, D.A. (2015). Waking State: Rapid Variations Modulate Neural and Behavioral Responses. Neuron 87, 1143–1161. 10.1016/J.NEURON.2015.09.012.

14. Lee, C.C.Y., Kheradpezhouh, E., Diamond, M.E., and Arabzadeh, E. (2020). State-Dependent Changes in Perception and Coding in the Mouse Somatosensory Cortex. Cell Rep. 32, 108197. 10.1016/j.celrep.2020.108197.

15. Akella, S., Ledochowitsch, P., Siegle, J.H., Belski, H., Denman, D.D., Buice, M.A., Durand, S., Koch, C., Olsen, S.R., and Jia, X. (2025). Deciphering neuronal variability across states reveals dynamic sensory encoding. Nat. Commun. 16, 1768. 10.1038/s41467-025-56733-w.

16. Zhu, Z., and Kuchibhotla, K. V. (2024). Performance errors during rodent learning reflect a dynamic choice strategy. Curr. Biol. 34, 2107–2117.e5. 10.1016/j.cub.2024.04.017.

17. Gallero-Salas, Y., Han, S., Sych, Y., Voigt, F.F., Laurenczy, B., Gilad, A., and Helmchen, F. (2020). Sensory and Behavioral Components of Neocortical Signal Flow in Discrimination Tasks with Short-Term Memory. Neuron 109, 1–14. 10.1016/j.neuron.2020.10.017.

18. Gilad, A., Gallero-Salas, Y., Groos, D., and Helmchen, F. (2018). Behavioral Strategy Determines Frontal or Posterior Location of Short-Term Memory in Neocortex. Neuron 99, 814–828.e7. 10.1016/j.neuron.2018.07.029.

19. Bolkan, S.S., Stone, I.R., Pinto, L., Ashwood, Z.C., Iravedra Garcia, J.M., Herman, A.L., Singh, P., Bandi, A., Cox, J., Zimmerman, C.A., et al. (2022). Opponent control of behavior by dorsomedial striatal pathways depends on task demands and internal state. Nat. Neurosci. 25, 345–357. 10.1038/s41593-022-01021-9.

20. Fries, P. (2005). A mechanism for cognitive dynamics: neuronal communication through neuronal coherence. Trends Cogn. Sci. 9, 474–480. 10.1016/J.TICS.2005.08.011.

21. Parto-Dezfouli, M., Vezoli, J., Bosman, C.A., and Fries, P. (2023). Enhanced behavioral performance through interareal gamma and beta synchronization. Cell Rep. 42, 113249. 10.1016/j.celrep.2023.113249.

22. Han, S., and Helmchen, F. (2024). Behavior-relevant top-down cross-modal predictions in mouse neocortex. Nat. Neurosci. 27, 298–308. 10.1038/s41593-023-01534-x.

23. Han, S., and Helmchen, F. (2025). Coordinated multi-level adaptations across neocortical areas during task learning. Nat. Commun. 16, 7719. 10.1038/s41467-025-62949-7.

24. Bandi, A.C., McCann, J.P., Carpenter, R.M., Lower, M.J., Long, J.N., and Runyan, C.A. (2025). Parietal cortex is causally required for state-dependent decisions. Cell Rep. 44, 116672. 10.1016/j.celrep.2025.116672.

25. Calhoun, A.J., Pillow, J.W., and Murthy, M. (2019). Unsupervised identification of the internal states that shape natural behavior. Nat. Neurosci. 22, 2040–2049. 10.1038/s41593-019-0533-x.

26. Mohammadi, Z., Ashwood, Z.C., and Pillow, J.W. (2025). Identifying the factors governing internal state switches during nonstationary sensory decision-making. Nat. Commun. 2025. 10.1038/s41467-025-66738-0.

27. Ashwood, Z.C., Roy, N.A., Stone, I.R., Urai, A.E., Churchland, A.K., Pouget, A., and Pillow, J.W. (2022). Mice alternate between discrete strategies during perceptual decision-making. Nat. Neurosci. 25, 201–212. 10.1038/s41593-021-01007-z.

28. Erlich, J.C., Brunton, B.W., Duan, C.A., Hanks, T.D., and Brody, C.D. (2015). Distinct effects of prefrontal and parietal cortex inactivations on an accumulation of evidence task in the rat. Elife 2015. 10.7554/ELIFE.05457.

29. Hwang, E.J., Dahlen, J.E., Mukundan, M., and Komiyama, T. (2017). History-based action selection bias in posterior parietal cortex. Nat. Commun. 8, 1–14. 10.1038/s41467-017-01356-z.

30. Semedo, J.D., Gokcen, E., Machens, C.K., Kohn, A., and Yu, B.M. (2020). Statistical methods for dissecting interactions between brain areas. Curr. Opin. Neurobiol. 65, 59–69. 10.1016/j.conb.2020.09.009.

31. Semedo, J.D., Zandvakili, A., Machens, C.K., Yu, B.M., and Kohn, A. (2019). Cortical Areas Interact through a Communication Subspace. Neuron 102, 249–259.e4. 10.1016/j.neuron.2019.01.026.

32. Nabbefeld, G., Abou Rachid, S., Lenzi, I., Cravo, E., Musall, S., and Kampa, B. (2025). Cortex-wide circuits for multisensory evidence accumulation and choice formation. bioRxiv, 19–22. 10.64898/2025.12.23.696238.

33. Pinto, L., Rajan, K., DePasquale, B., Thiberge, S.Y., Tank, D.W., and Brody, C.D. (2019). Task-Dependent Changes in the Large-Scale Dynamics and Necessity of Cortical Regions. Neuron 104, 810–824.e9. 10.1016/j.neuron.2019.08.025.

34. MacDowell, C.J., and Buschman, T.J. (2020). Low-Dimensional Spatiotemporal Dynamics Underlie Cortex-wide Neural Activity. Curr. Biol. 30, 2665–2680.e8. 10.1016/j.cub.2020.04.090.

35. Lee, J.J., Krumin, M., Harris, K.D., and Carandini, M. Task specificity in mouse parietal cortex. 10.1101/2020.12.18.423543.

36. Orsolic, I., Rio, M., Mrsic-Flogel, T.D., and Znamenskiy, P. (2021). Mesoscale cortical dynamics reflect the interaction of sensory evidence and temporal expectation during perceptual decision-making. Neuron 109, 1861–1875.e10. 10.1016/j.neuron.2021.03.031.

37. Hira, R., Townsend, L.B., Smith, I.T., Yu, C.-H., Stirman, J.N., Yu, Y., and Smith, S.L. (2026). Mesoscale functional architecture in medial posterior parietal cortex. Elife 14. 10.7554/eLife.105213.3.

38. Harvey, C.D., Coen, P., and Tank, D.W. (2012). Choice-specific sequences in parietal cortex during a virtual-navigation decision task. Nature 484, 62–68. 10.1038/nature10918.

39. Kira, S., Safaai, H., Morcos, A.S., Panzeri, S., and Harvey, C.D. (2023). A distributed and efficient population code of mixed selectivity neurons for flexible navigation decisions. Nat. Commun. 14, 2121. 10.1038/s41467-023-37804-2.

40. Guyoton, M., Matteucci, G., Foucher, C.G., Getz, M.P., Gjorgjieva, J., and El-Boustani, S. (2025). Cortical circuits for cross-modal generalization. Nat. Commun. 16, 4230. 10.1038/s41467-025-59342-9.

41. Pho, G.N., Goard, M.J., Woodson, J., Crawford, B., and Sur, M. (2018). Task-dependent representations of stimulus and choice in mouse parietal cortex. Nat. Commun. 9. 10.1038/s41467-018-05012-y.

42. Zhong, L., Zhang, Y., Duan, C.A., Deng, J., Pan, J., and Xu, N. long (2019). Causal contributions of parietal cortex to perceptual decision-making during stimulus categorization. Nat. Neurosci. 22, 963–973. 10.1038/s41593-019-0383-6.

43. Freedman, D.J., and Ibos, G. (2018). An Integrative Framework for Sensory, Motor, and Cognitive Functions of the Posterior Parietal Cortex. Neuron 97, 1219–1234. 10.1016/j.neuron.2018.01.044.

44. Zhao, J., An, B., Cheng, N., Dong, Q., Wang, C., Correspondence, C., and Chen, X. (2025). Gravity-dependent choice of frame of reference in the posterior parietal cortex. Curr. Biol. 0. 10.1016/J.CUB.2025.08.048.

45. Morcos, A.S., and Harvey, C.D. (2016). History-dependent variability in population dynamics during evidence accumulation in cortex. Nat. Neurosci. 19, 1672–1681. 10.1038/nn.4403.

46. Choi, I., and Lee, S.-H. (2025). Locomotion-dependent auditory gating to the parietal cortex guides multisensory decisions. Nat. Commun. 16, 2308. 10.1038/s41467-025-57347-y.

47. Gilad, A., and Helmchen, F. (2020). Spatiotemporal refinement of signal flow through association cortex during learning. Nat. Commun. 11, 1–14. 10.1038/s41467-020-15534-z.

48. Zhu, Q., Wang, Y., Huang, Y., Yang, X., and Guo, Z. V. (2026). Posterior parietal cortex guides sensorimotor associative learning by linking sensation to distal action. bioRxiv, 2026.04.19.719425. 10.64898/2026.04.19.719425.

49. Arlt, C., Barroso-Luque, R., Kira, S., Bruno, C.A., Xia, N., Chettih, S.N., Soares, S., Pettit, N.L., and Harvey, C.D. (2022). Cognitive experience alters cortical involvement in goal-directed navigation. Elife 11. 10.7554/ELIFE.76051.

50. Gilissen, S.R.J., Farrow, K., Bonin, V., and Arckens, L. (2021). Reconsidering the Border between the Visual and Posterior Parietal Cortex of Mice. Cereb. Cortex 31, 1675–1692. 10.1093/CERCOR/BHAA318.

51. Tseng, S.-Y., Chettih, S.N., Arlt, C., Barroso-Luque, R., and Harvey, C.D. (2022). Shared and specialized coding across posterior cortical areas for dynamic navigation decisions. Neuron 110, 2484–2502.e16. 10.1016/j.neuron.2022.05.012.

52. Yu, Y., Stirman, J.N., Dorsett, C.R., and Smith, S.L. (2026). Visual information is broadcast among cortical areas in discrete channels. Elife 13, 469114. 10.7554/eLife.97848.3.

53. Orlandi, J.G., Abdolrahmani, M., Aoki, R., Lyamzin, D.R., and Benucci, A. (2023). Distributed context-dependent choice information in mouse posterior cortex. Nat. Commun. 14, 192. 10.1038/s41467-023-35824-6.

54. Rowland, J.M., van der Plas, T.L., Loidolt, M., Lees, R.M., Keeling, J., Dehning, J., Akam, T., Priesemann, V., and Packer, A.M. (2023). Propagation of activity through the cortical hierarchy and perception are determined by neural variability. Nat. Neurosci. 26, 1584–1594. 10.1038/s41593-023-01413-5.

55. Ebrahimi, S., Lecoq, J., Rumyantsev, O., Tasci, T., Zhang, Y., Irimia, C., Li, J., Ganguli, S., and Schnitzer, M.J. (2022). Emergent reliability in sensory cortical coding and inter-area communication. Nature 605, 713–721. 10.1038/s41586-022-04724-y.

56. Marrero, K., Aruljothi, K., Zareian, B., Gao, C., Zhang, Z., and Zagha, E. (2022). Global, Low-Amplitude Cortical State Predicts Response Outcomes in a Selective Detection Task in Mice. Cereb. Cortex 32, 2037–2053. 10.1093/cercor/bhab339.

57. Reimer, J., Froudarakis, E., Cadwell, C.R., Yatsenko, D., Denfield, G.H., and Tolias, A.S. (2014). Pupil Fluctuations Track Fast Switching of Cortical States during Quiet Wakefulness. Neuron 84, 355–362. 10.1016/j.neuron.2014.09.033.

58. Picciotto, M.R., Higley, M.J., and Mineur, Y.S. (2012). Acetylcholine as a Neuromodulator: Cholinergic Signaling Shapes Nervous System Function and Behavior. Neuron 76, 116–129. 10.1016/j.neuron.2012.08.036.

59. Collins, L., Francis, J., Emanuel, B., and McCormick, D.A. (2023). Cholinergic and noradrenergic axonal activity contains a behavioral-state signal that is coordinated across the dorsal cortex. Elife 12. 10.7554/ELIFE.81826.

60. Suzuki, M., and Larkum, M.E. (2020). General Anesthesia Decouples Cortical Pyramidal Neurons. Cell 180, 666–676.e13. 10.1016/j.cell.2020.01.024.

61. Valente, M., Pica, G., Bondanelli, G., Moroni, M., Runyan, C.A., Morcos, A.S., Harvey, C.D., and Panzeri, S. (2021). Correlations enhance the behavioral readout of neural population activity in association cortex. Nat. Neurosci. 24, 975–986. 10.1038/s41593-021-00845-1.

62. Chen, J.L., Voigt, F.F., Javadzadeh, M., Krueppel, R., and Helmchen, F. (2016). Long-range population dynamics of anatomically defined neocortical networks. Elife 5. 10.7554/eLife.14679.

63. Pachitariu, M., Stringer, C., Dipoppa, M., Schröder, S., Rossi, L.F., Dalgleish, H., Carandini, M., and Harris, K.D. (2016). Suite2p: beyond 10,000 neurons with standard two-photon microscopy. bioRxiv. 10.1101/061507.

